# The “osteostat”: a theory of bone mechanosensing and setpoint adaptation based on osteocytes

**DOI:** 10.64898/2026.06.23.734120

**Authors:** Yves Pauchard, Pascal R Buenzli

**Affiliations:** Department of Electrical and Software Engineering, University of Calgary, Calgary, Canada; School of Mathematical Sciences, Queensland University of Technology (QUT), Brisbane, Australia

**Keywords:** bone tissue regulation, mechanical adaptation, osteocyte, mechanostat, bone remodelling, lazy zone

## Abstract

The osteocyte network in bone is believed to play an important role for how bone tissues sense and respond to mechanical stimulation. Yet, bone adaptation to mechanical loads is often conceptualised as a simple response to mechanical stimuli, such as Wolff’s law, which is based on mechanical variables only and takes no account of the cellular basis of mechanosensation. Wolff’s law presumes the existence of a reference mechanical stimulus, the mechanical setpoint, above which bone is added , and under which bone is removed. In this paper, we develop a theory of bone tissue sensing and adaptation based on osteocytes to provide new understanding of the role played by osteocyte signals in mechanical adaptation. In this theory, the mechanical setpoint of Frost’s mechanostat is explicitly embodied as osteocyte properties involved in mechanotransduction. The mechanical setpoint is allowed to adapt due to the replacement of osteocytes during remodelling, making the setpoint space and time dependent. We propose a mathematical model to implement this new theory of bone adapation and present numerical simulations of this model to explore how mechanobiological response curves (effective Wolff’s laws) are modulated by setpoint adaptation during remodelling. By accounting for varying osteocyte populations within bone tissue, we explore bone adaptation under osteocyte disruptions, which is particularly relevant to age-related bone loss. Our model suggests that biological disruptions of remodelling balance cannot always be compensated by mechanical feedback, and that setpoint adaptation during remodelling may have significant observable consequences, such as hysteresis in bone response signatures that resemble lazy zones.

## 1 Introduction

Bone tissue is equipped with sensing and repair mechanisms that enable shape adaptation and the repair of micro-cracks, so bone can be light and mechanically robust [1]. The adaptation of bone shape and architecture to mechanical load has been formalised by Frost as the “mechanostat” theory [2, 3]. This theory describes how bone changes with respect to a mechanical reference state called the *mechanical setpoint*. The mechanostat is similar to how a thermostat senses and regulates a room’s temperature to a pre-defined value. Regions of bone experiencing mechanical stress above the setpoint are consolidated to reduce mechanical stress, while regions experiencing mechanical stress below the setpoint are removed to reduce mass and increase mechanical stress toward the setpoint.

A central question in the mechanostat theory of bone is how a mechanical setpoint emerges from the biological underpinning of bone. Frost introduced the mechanical setpoint as a quantity that may be influenced by several regulatory factors, to help explain the regulation of bone tissues in health and disease [2]. Many following studies expanded on the nature of the setpoint to emphasise it may depend on skeletal site and a variety of local and systemic biological factors [3–8]. To explain reduced bone responses to sustained mechanical stimulus, Turner [9] also proposed that the setpoint evolves over short times due to cellular accommodation to new loads. The time-dependence of the setpoint is evidenced by the load-history dependence of mechanical adaptation [8–11]. These observations clearly emphasise that bone structure is not merely in one-to-one correspondence with purely mechanical variables, it is also the result of how mechanics is sensed and transduced biologically by dynamic cellular processes. In other words, the setpoint is not a mechanical property, it is a property of cellular signals and actions. The mechanostat is well established as a conceptual theory, but it has remained difficult to experimentally test some of its features directly, including how it depends on cellular processes. A number of direct experimental observations support the mechanostat theory as a structural response of bone to mechanical loads. Bone response to mechanical overstimulation in animal models have provided experimental insights into Wolff’s laws of bone adaptation (response curves to mechanical stimulus), and demonstrated that these responses vary with age [8, 10, 12–18]. Bone response curves to mechanical stimulus have also been estimated in humans [19]. However, it remains unknown how the mechanical setpoint itself relates to material, cellular, and molecular variables and their change in space and time. One of the challenges in investigating a mechanical setpoint experimentally is to distinguish bone formation and resorption associated with normal physiological conditions such as homeostatic remodelling, from bone formation and resorption induced by mechanical stimulation, such as overload or underload. While the mechanical setpoint has in principle a clear practical definition based on bone responses, the complexity of bone adaptation and the difficulty of measuring bone response curves mean that the mechanical setpoint has remained an elusive quantity with no clearly identified biological underpinning.

In this paper, we put forward the hypothesis that the cellular processes from which an effective setpoint emerges in mechanical bone adaptation can be attributed in whole or in part to the osteocyte network embedded in bone tissues. This new theory of bone mechanobiology extends Frost’s mechanostat theory by explicitly including osteocytes as the cells representing the mechanical setpoint and regulating mechanical adaptation, which we therefore propose to call the “osteostat” theory of mechanical adaptation. Osteocytes, long thought to be quiescent cells within hard bone tissue, have emerged as central players for the mechanical adaptation of bone tissues [8, 20–27]. Osteocytes form a dense cellular network embedded within bone tissue [28]. This allows osteocytes to sense mechanical deformations of the tissue, and to respond to these deformations by propagating signalling molecules to the bone surface, where they may induce osteogenic or osteolytic responses by stimulating the generation of osteoblasts or osteoclasts [27, 29–33]. These osteogenic and osteolytic responses, in turn, change bone mass and micro-architecture to bring mechanical deformations of the tissue closer to the setpoint. While this is a common view of how bones adapt their mass and structure to mechanical loads, many questions about the implications of the osteocytic control of bone tissue regulation remain unexplored, and an osteocyte-centred theory of bone tissue regulation is currently lacking.

The osteocyte network is a dynamic entity subject to modelling and remodelling processes. Osteocytes may be created, replaced, and they may be removed by bone resorption or undergo apoptosis. All these processes are expected to influence the mechanotransduction of bone tissue mechanics by which mechanical deformations of the tissue are detected and transduced into biological signalling molecules triggering bone formation and bone resorption [33]. The creation and removal of osteocytes during bone formation and bone resorption may therefore change response mechanisms in space and time and provide a long-term adaptation of the setpoint in addition to the short-term cellular accommodation proposed by Turner [9].

The idea that the setpoint may be encoded in osteocyte properties was first proposed in our previous work [34]. In this work, the osteocyte population was considered to be static, i.e., osteocyte density within bone tissue was assumed to be constant and there was no distinction between lacunar density and osteocyte density. Here we extend this model with a dynamic osteocyte population to investigate how mechanical adaptation is influenced by disruptions of osteocytes that may occur with age or disease. With age, osteocyte density may decrease due to lower lacunar densities [35–38], lower lacunar occupancy [35, 37, 39–41], and increased osteocyte apoptosis [42–46]. Further influences on osteocytes are known, in particular, mechanical unloading is shown to increase apoptosis in animal models [47] and microgravity decreases osteocyte populations [48,49]. Osteocyte apoptosis is linked to the development of resorption-related bone diseases including osteoporosis and fragility fractures [50]. The aim of the current study is to use the proposed osteostat theory to find qualitative features arising from assumed osteocyte-specific regulation that may be observable experimentally. To ultimately help design new experiments that investigate these processes, it is useful to conceptualise the regulatory role of osteocytes into a mathematical model and to explore the implication of this model in a number of concrete, albeit idealised situations, using numerical simulations. As such, the proposed model is not to be interpreted as a quantitative model. We explore the implication of a changing osteocyte population for bone sensing and adaptation in health and in aging or disease, illustrated by means of a mathematical model at the tissue scale. The model follows temporal changes in bone cell populations and in bone mass, and represents these changes as effective Wolff-law type bone response curves.

## 2 The “osteostat”: a theory of bone tissue sensing and regulation

In this section, we present the *osteostat* as a conceptual model of the role played by osteocytes in the mechanical adaptation of bone tissues. This conceptual model is based on biological assumptions about how osteocytes sense tissue mechanics and how they regulate osteoblasts and osteoclasts. In Section 3, we transcribe this conceptual model into a mathematical model to explore the implications of these biological assumptions quantitatively.

### Assumptions of the osteostat

The main assumptions of our osteocyte-based theory of the functional adaptation of bone are as follows (Figure 1):

**Figure 1.**
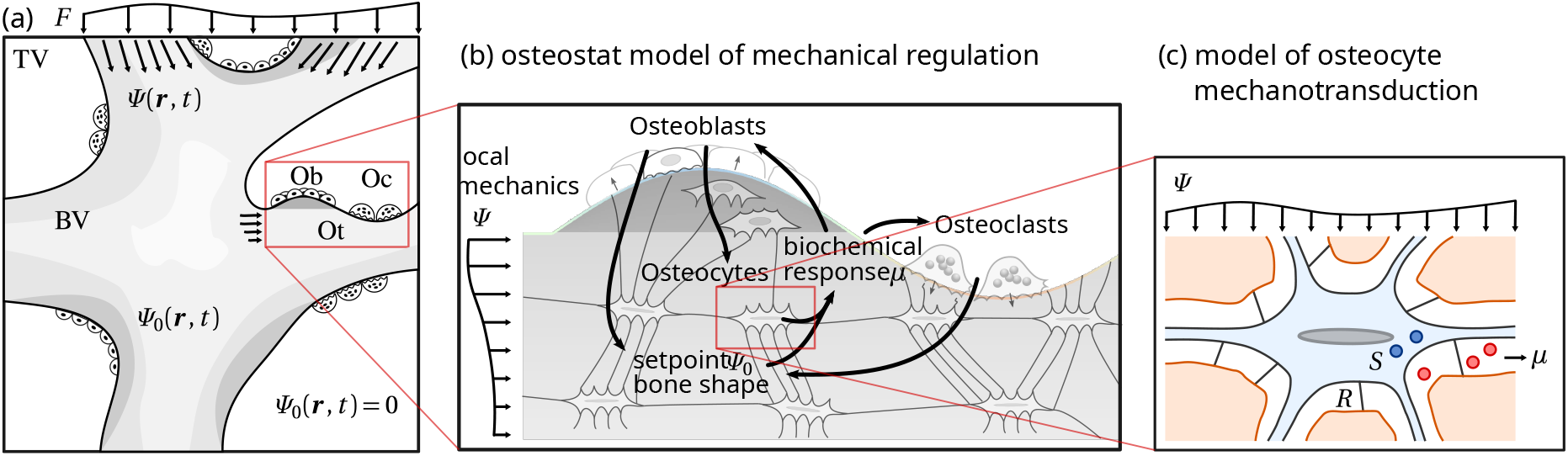
Osteostat model. (a) Mechanical loads applied to a reference tissue volume (TV) generate space and time dependent mechanical stimulus *Ψ* (***r*** , *t*) in the bone volume (BV). The mechanical setpoint *Ψ*_0_(***r*** , *t*) in BV is a bone material property related to osteocytes; (b) Osteocytes generate a mechanotransduced signal *µ* in response to the comparison between *Ψ* (***r*** , *t*) and the setpoint *Ψ*_0_(***r*** , *t*). The signal *µ* propagates to the bone surface, where it initiates osteoblast or osteoclast formation. Osteoblasts and osteoclasts change bone mass and therefore local mechanics *Ψ* (***r*** , *t*). In addition, new osteocytes are generated during bone formation, updating the value of the mechanical setpoint *Ψ*_0_(***r*** , *t*); (c) Osteocytes transduce the local mechanical stimulus *Ψ* through tethering elements *R* , triggering intracellular chemical pathways *S* and the release of a mechanotransduced signal *µ*.

H1 The osteocyte network is able to sense local tissue mechanics (Figure 1a). Osteocytes respond to mechanical stimulus by emitting osteogenic signals when experiencing overload, and osteolytic signals when experiencing underload. These signals propagate through the osteocyte network or lacuno-canalicular network to the bone surface, where they induce osteoblastogenesis and osteoclastogenesis (Figure 1b).

H2 Whether osteocytes experience overload or underload depends on the local tissue mechanical environment, which we represent here by a variable *Ψ* , as well as how *Ψ* is perceived and transduced by the osteocyte network (Figure 1a–c). Our main assumption is that intrinsic properties of osteocytes involved in mechanotransduction, such as the osteocyte network architecture, osteocyte density, and genotypic and phenotypic cell properties, define and materialise a reference mechanical state *Ψ*_0_ that acts as a mechanical setpoint. The space and time dependence of the setpoint is due to local variations of these intrinsic osteocyte properties, that can change over long time scales (months–years) when new osteocytes are generated, or over short timescales (hours–days) due to cellular accommodation.

H3 Bone mass changes by the action of osteoblasts and osteoclasts. Bone resorption removes old osteocytes and their associated setpoint. Bone formation generates osteocytes with new mechanotransduction properties, that may be influenced by the current mechanical environment. It associates newly formed bone with a new setpoint value.

Hypothesis 1 is based on our current understanding of osteocyte signalling in bone. A large body of literature is devoted to the question of which mechanical stimuli may be sensed by osteocytes [7, 8, 15, 51–54], and how signals may propagate through the network [30–33, 54–58]. These considerations are all consistent with the assumptions of Hypothesis 1. The osteostat does not rely on a particular mechanosensation mechanism nor a particular choice of mechanical variable. In the mathematical model, we will assume the mechanical stimulus *Ψ* to be the strain energy density for illustration purposes, but other quantities could be considered within our framework.

Hypothesis 2 assumes that when new osteocytes are created, they make up a new functional syncytium that is homeostatic for the mechanical and physiological conditions that prevail at the time and location of their creation [59]. Experimentally, the osteocyte network has been shown to possess different morphologies in different skeletal sites [27, 60–64]. How network topology, osteocyte morphology and other intrinsic properties of the osteocyte network are generated in different bone tissues (lamellar bone, woven bone) and in different skeletal sites (cortical bone, trabecular bone, endosteum, periosteum, neutral axis), depends on mechanical and biological factors at the time of new bone formation, including physiological and microenvironmental conditions such as repair and pathological responses. While osteocyte shape in woven bone may be more predominantly influenced by biological factors, osteocyte shape and orientation in lamellar bone may be more predominantly influenced by mechanical factors [61]. It is reasonable to expect that local differences in intrinsic properties of the osteocyte network generate long-term influences on mechanosensitivity and participate in determining a local mechanical reference state (setpoint) [34, 61, 65–67]. The extent to which newly generated osteocytes are adapted to the mechanical environment that they experience at the time and location of their creation is currently unknown. In the mathematical model, we will assume that osteocytes are created entirely adapted as a working hypothesis, with the aim to explore experimentally observable consequences of this hypothesis, that may be tested in future experiments. Hypotheses 3 is based on a common view of the regulatory role of osteocytes for bone mass changes, which involves the action of osteoblasts and osteoclasts at the bone surface [21, 68–71]. While the mathematical model proposes specific mechanisms for encoding the mechanical setpoint in osteocyte-specific properties for illustration purposes, we also show that this detail is not important for the qualitative behaviour of the model. The mechanical setpoint *Ψ*_0_ can be viewed as a bone tissue material property which is evolved like any other material property during bone remodelling, with old values of *Ψ*_0_ being replaced by the mechanical stimulus *Ψ* that prevails at the time of bone formation, with the assumption that newly formed bone is mechanically adapted at the time of formation. However, the detail of which specific property encodes the setpoint could be biologically relevant to understand what may impact changes to the setpoint during the lifetime of a newly formed region of bone.

A number of physiological conditions may influence osteocytes and their regulatory function for bone adaptation and repair. The density of osteocytes is first determined during bone formation. This density remains unchanged by bone resorption so long as resorption indiscriminately removes bone of any osteocyte density on average (see Figure 2). However, the density of osteocytes in BV may change when osteocytes die by apoptosis. Several factors are known to influence osteocyte appoptosis, including aging, estrogen deficiency, and glucocorticoid-induced apoptosis [46]. Furthermore, it has been shown that osteocytes undergo apoptosis when mechanical load is reduced [72], so that mechanical loading can influence apoptosis rate and osteocyte density. As osteocytes age or undergo apoptosis, the osteocyte network becomes disrupted, which significantly impairs the detection of mechanical signals and leads to increased bone fragility [73]. Consequently, the accumulation of dead osteocytes may cause the skeleton to no longer effectively repair microdamage. Dead osteocytes leave behind empty lacunae that are often mineralised, leading to so-called “micropetrosis” [74].

**Figure 2.**
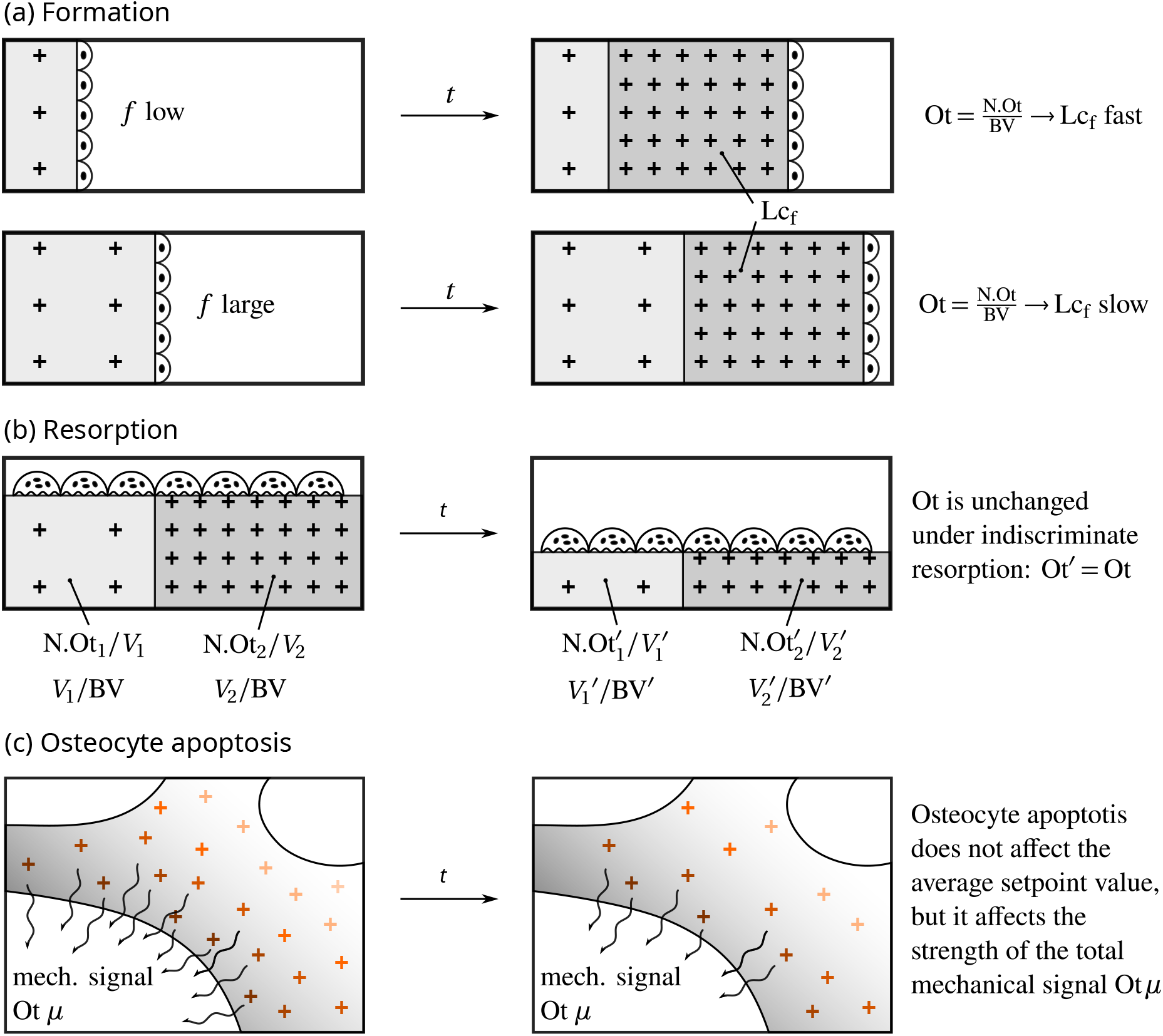
(a) Evolution of osteocyte density under bone formation. When bone volume fraction *f* is low, the average osteocyte density Ot (crosses) converges more rapidly to the lacunar density Lc_f_ of newly formed bone than when *f* is large; (b) Evolution of osteocyte density under bone resorption. If resorption is indiscriminate and targets any bone region regardless of osteocyte density, then the average osteocyte density is unchanged: resorption does not affect the local osteocyte densities, so that 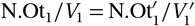 and 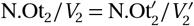, and it does not affect the fractions of bone volume in which these densities reside, i.e 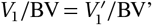’ and 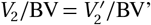. Therefore, Ot = (N.Ot_1_*/V*_1_)(*V*_1_*/*BV) +(N.Ot_2_*/V*_2_)(*V*_2_*/*BV) = Ot^*′*^ (see also Appendix A); (c) Influence of osteocyte apoptosis for mechanical adaptation. The total mechanical signal received by osteoclasts and osteoblasts is proportional to Ot *µ*, and therefore affected by osteocyte death. In contrast, the mechanical setpoint *Ψ*_0_ (colour of the crosses) is unaffected by osteocyte death, since this intensive quantity represents average mechanical sensitivies of live osteocytes.

Despite the many factors influencing osteocyte death, these cells are considered to have a long biological lifespan. In the absence of remodeling, these cells can live for decades. In dense cortical bone, where remodelling rates are low, on the order of 4%/year [75, Table 1], they may persist for over 25 years [21]. However, their lifespan in high-turnover trabecular bone with remodelling rates around 28%/year [75, Table 1] is significantly shorter, on the order of 3–4 years.

**Table 1.** Symbols used in the mathematical model and their correspondence with standard histomorphometric nomenclature [76]. Conversion from volume referent BV to TV is obtained by multiplication by the bone volume fraction f . Model parameters and their values are listed in Table 2 in Appendix A.

| Mathematical notation | Histomorphometric notation | Description | Volume referent |
| --- | --- | --- | --- |
| $f$ | BV/TV | bone volume fraction [–] | TV |
| $S_V$ | BS/TV | specific surface [$\text{mm}^{-1}$] | TV |
| Ob | N.Ob/TV | osteoblast density [$\text{mm}^{-3}$] | TV |
| Oc | N.Oc/TV | osteoclast density [$\text{mm}^{-3}$] | TV |
| Ot | N.Ot/BV | osteocyte density [$\text{mm}^{-3}$] | BV |
| Lc | N.Lc/BV | lacunar density [$\text{mm}^{-3}$] | BV |
| $\Psi$ | – | strain energy density [GPa] | BV |
| $\Psi_0$ | – | mechanical setpoint [GPa] | BV |
| $\mu$ | – | transduced mechanical signal [–] | BV |
| $k_f$ | – | secretory rate per Ob [$\mu\text{m}^3/\text{day}$] | – |
| $k_r$ | – | resorption rate per Oc [$\mu\text{m}^3/\text{day}$] | – |
| $\frac{1}{f} k_f \text{Ob}$ | BFR = BFR/BV | bone formation rate [$\text{day}^{-1}$] | BV |
| $\frac{1}{f} k_r \text{Oc}$ | BRs.R = BRs.R/BV | bone resorption rate [$\text{day}^{-1}$] | BV |

## 3. Mathematical model

We now formalise our conceptual model of the osteostat presented in Section 2 into a mathematical model. We generalise our previous mathematical model of osteoblast and osteoclast populations and setpoint adaptation [34] by accounting for the influence of a dynamic population of osteocytes within BV, which affects the strength of the mechanical signal received by osteoblasts and osteoclasts. Here, we present a summary of key equations of the mathematical model, followed by a brief discussion of their structure and assumptions. The reader is referred to Appendix A for a more detailed presentation. The mathematical model introduces a number of mathematically convenient notations; Table 1 below lists these notations and their correspondence with standardised histomorphometric notation [76]. Table 2 in Appendix A lists model parameters and their values.

**Table 2.**
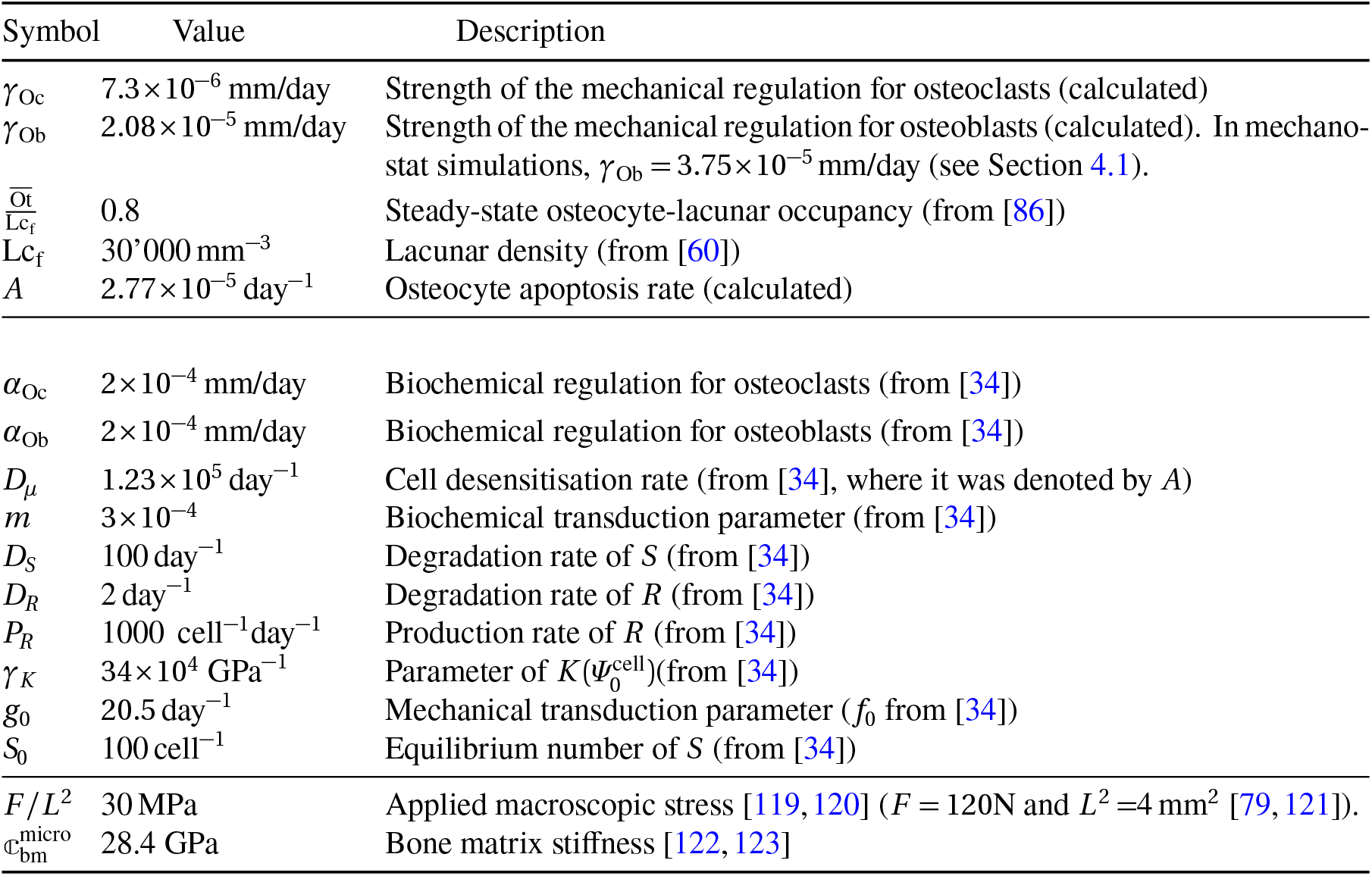
Model parameters.

We consider the time evolution of spatially averaged quantities in a fixed region of bone tissue TV of the order of 2–8 mm^3^. To formulate our model of cell populations, we assume that bone architecture in this region can be characterised by two microstructural parameters, bone volume fraction *f* = BV*/*TV, and the specific surface *S*_V_ = BS*/*TV. Experimental measurements show that the specific surface *S*_V_ is largely determined by bone volume fraction, so that bone volume fraction alone is a good indicator of microarchitecture [77–80]. The evolution of bone microarchitecture is therefore represented well by the evolution of bone volume fraction within TV (Figure 1a). At high values of *f* , for example, experimental measurements of *S*_V_ are consistent with microstructures made of cylindrical pores of varius radii [79]. The time rate of change of *f* is given by

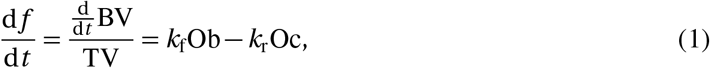

where Ob = N.Ob/TV and Oc = N.Oc/TV are the densities of osteoblasts and osteoclasts in TV, *k*_f_ is the volume of bone formed per unit time per osteoblast, and *k*_r_ is the volume of bone resorbed per unit time per osteoclast. We assume that the populations of osteoblasts and osteoclasts is made up of a physiological baseline population responsible for bone remodelling, denoted by a superscript (b), and a mechanics-induced population generated in response to mechanical adaptation, denoted by a superscript (m). Each of these cell populations is assumed to be proportional to the vascular space 1 *−f* , which provides precursor cells, and to the amount of bone surface via the specific surface *S*_V_(*f*) = BS*/*TV, which is an experimentally determined function of the bone volume fraction *f* [77, 78, 81]:

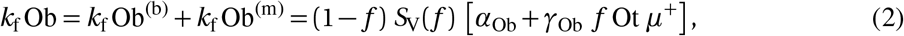

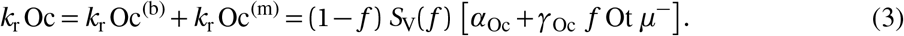

In Eqs (2)–(3), *α*_Ob_ and *α*_Oc_ represent the physiological baseline contribution. The mechanics-induced contributions, weighted by parameters *γ*_Ob_ and *γ*_Oc_, are assumed to be proportional to the total number of osteocytes in TV, N.Ot/TV = *f* Ot, and to the positive part *µ*^+^ = max(*µ*, 0) *>* 0 or negative part *µ*^*−*^ = *−*min(*µ*, 0) *>* 0 of a mechanical stimulus signal *µ*. These positive and negative parts are such that *µ >* 0 generates additional osteoblasts in proportion to |*µ*, and *µ <* 0 generates additional osteoclasts in proportion to *µ*| . The parameters *γ*_Ob_ and *γ*_Oc_ determine the extent to which mechanical overloading and underloading modulate osteoblast and osteoclast populations. These parameters can be calibrated against experimental bone responses to mechanical stimulation, see Section 4.1. Several options can be devised for an appropriate mechanical stimulus |*µ*|. In the simplest case, the mechanical stimulus *µ* can be defined as

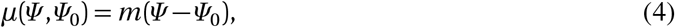

where *Ψ*_0_ is the mechanical setpoint and *m* is a scaling factor. A more sophisticated version of *µ* is presented in Appendix A, based on a model of osteocyte mechanotransduction.

The mechanical setpoint itself evolves due to bone formation and resorption according to

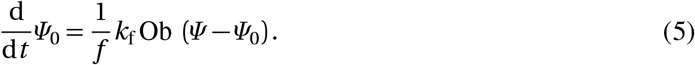

Equation (5) represents Hypothesis H2 of the osteostat (see Section 2). Further details about how the setpoint evolution can be related to evolving osteocyte properties are presented in Appendix A and Appendix B. Resorption only affects the evolution of the setpoint implicitly in Eq. (5), via changes in *f* , see the discussion below. Finally, the density of osteocytes Ot within BV evolves due to bone formation, bone resorption, and cell death (such as apoptosis) according to

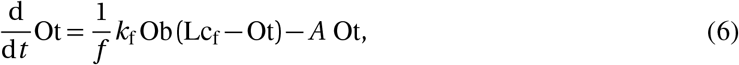

where Lc_f_ = N.Lc/BV is the density of osteocyte lacunae generated in newly formed bone, and *A* is a cell death rate parameter [82, 83].

Osteocyte lifespan is hard to measure experimentally, but Equation (6) provides a way to estimate osteocyte lifespan from the knowledge of remodelling rate and osteocyte occupancy Ot*/*Lc_f_. Our model includes two mechanisms by which osteocytes may be eliminated: cell death, which results in empty lacunae, and bone resorption, which removes both osteocytes and lacunae. Osteocyte lifespan can be measured as the effective half-life *T*_1*/*2_, which represents the time after which the survival probability of an osteocyte is reduced by half. In a steady state with no overall bone loss nor gain (*k*_f_Ob = *k*_r_Oc), the effective half-life *T*_1*/*2_ of an osteocyte due to apoptosis and removal by remodelling, is given by (see Appendix A):

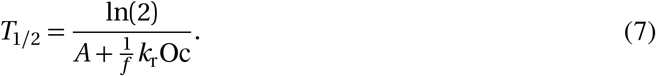

Equations (1)–(6) form a system of ordinary differential equations (ODEs) that can be solved numerically using standard ODE solvers, see Section A.1 for more detail. We now conclude this section with a brief discussion of the structure and assumptions of the mathematical model:

1. The assumption that the mechanics-induced population of osteoblasts and osteoclasts in Eqs (2)– (3) is proportional to Ot *µ* can be justified by the high connectivity of the osteocyte network, which suggests that signals may be integrated over volumes containing several bone structural units [28, 82]. This assumption means that the more osteocytes there are in TV, the stronger the bone response, and conversely, the fewer osteocytes there are in TV, the weaker the bone response. In contrast, the mechanical stimulus *µ* defined in Eq. (4) is an averaged quantity that does not scale in proportion to bone volume.
2. Equation (4) is similar to Wolff’s law, in that it generates an osteogenic response if *Ψ > Ψ*_0_ and an osteolytic response if *Ψ < Ψ*_0_. However, it differs from such a simple response law by proposing a material embodiment of the setpoint *Ψ*_0_ associated with osteocyte-specific properties. This material embodiment of the setpoint allows us to prescribe precisely how *Ψ*_0_ evolves due to changes in these osteocyte properties in Eq. (5) (see also Appendix A). The mechanical stimulus *µ* represents a signalling compound emitted by osteocytes through mechanotransduction pathways in response to the discrepancy between *Ψ* and the setpoint *Ψ*_0_. For simplicity, we do not include short-term cell desensitisation mechanisms proposed by Turner [9] in the mathematical model presented here, and only focus on long-term influences on *µ* due to bone resorption and formation. The reader is referred to Ref. [34] for an osteocyte transduction model of *µ* that includes both short-term cell accommodation and long-term adaptation due to bone remodelling.
3. A major assumption of the mathematical model is that when new osteocytes are generated during bone formation, they are born adapted to their current mechanical environment. This assumption is reflected in Eq. (5) by the average setpoint *Ψ*_0_ always evolving toward the current strain energy density *Ψ* during bone formation: if *Ψ > Ψ*_0_, then the rate of change of *Ψ*_0_ is positive and *Ψ*_0_ increases toward *Ψ* ; if *Ψ < Ψ*_0_, then the rate of change of *Ψ*_0_ is negative and *Ψ*_0_ decreases toward *Ψ* . The evolution equation of *Ψ*_0_ arises from spatially averaging a spatio-temporal balance equation of osteocyte properties under bone formation and resorption, see Appendix B.
4. The rate at which *Ψ*_0_ tends to new setpoint values *Ψ* in Eq. (5) and at which Ot tends to new osteocyte densities Lc_f_ in Eq. (6) due to new bone formation is clearly proportional to the bone formation rate *k*_f_Ob. Figure 2a illustrates why these rates are also inversely proportional to bone volume fraction *f* . Figure 2b illustrates the fact that resorption does not affect the evolution of Ot because Ot in the mathematical model represents an average density of osteocytes with BV as volume referent, and because resorption is assumed to indiscriminately remove bone within TV that contains densities of osteocytes equal to the current average density within BV. Similar reasonings justify why Eq. (5) does not explicitly depend on bone resorption. These properties can be shown mathematically [83].
5. While the setpoint *Ψ*_0_ is a material embodiment of osteocyte-specific properties, its evolution in Eq. (5) is unaffected by osteocyte death. This represents the assumption that osteocytes with new mechanical sensitivities cannot regenerate in empty lacunae. However, osteocyte death influences osteocyte density Ot, and therefore the total strength of mechanical response within TV, see point 1 above and Figure 2c.

## 4 Model simulation experiments

In this section, we first calibrate the mathematical model outlined above, and then use the calibrated model to explore some its properties under changes to mechanical loading and under osteocyte disruptions.

### 4.1 Model calibration

The mathematical model contains a number of parameters, summarised in Table 2 in Appendix A. Most of these parameters are taken directly from our previous study [34, Table A.1], which were based on values from the biological literature, or estimated by calibration to astronaut data [84, 85]. New parameters introduced in our present study are estimated as follows.

Osteocyte apoptosis rate *A* is estimated based on experimental data on osteocyte occupancy and remodelling rate. In a steady state with no bone gain nor loss within TV, a constant apoptosis rate *A*, and a baseline of bone turnover creating new osteocytes, Eq. (6) shows that the average osteocyte occupancy Ot*/*Lc_f_ has the steady-state value

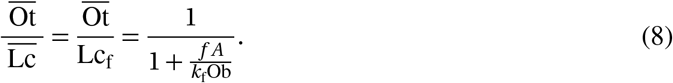

The rate parameter *A* can be estimated from Eq. (8), assuming an osteocyte occupancy 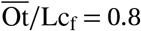 (80%) [86] in bone of volume fraction *f* = 0.85 remodelling at rate *α* = *α*_Ob_ = *α*_Oc_ = 2 *×* 10^*−*4^ mm/day, giving *A* = 2.77 *×* 10^*−*5^ day^*−*1^. Our numerical simulations also assume an average lacunar density Lc_f_ = 30000 */*mm^3^ [60], so that in a steady state with 80% lacunar occupancy, 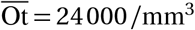.

The mathematical model in Ref. [34] had constant mechanical regulation parameters *β*_Ob_, *β*_Oc_ which are now extended to time-varying quantities *γ*_Ob_Ot(*t*) and *γ*_Oc_Ot(*t*) in Eqs (2), (3), respectively. The new mechanical regulation parameters *γ*_Ob_, *γ*_Oc_ are calibrated based on long-time unloading and reloading experiments presented in Section 4.2. We set 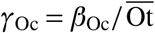 to ensure that the mechanical regulation right after the onset of unloading, where 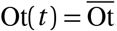, matches the response of the previous model [34]. However, instead of setting the value of *γ*_Ob_ to 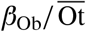, we set 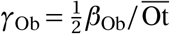 because in our unloading-reloading experiment (Section 4.2), the value of Ot has increased by the time we reload. Without the extra reduction of *γ*_Ob_ by half compared to *β*_Ob_*/* Ot, the response after reloading would be too large compared to the previous model [34].

Figure 3 compares numerical simulations of the model with experimental data from astronauts undertaking long space missions (4–6 months) before returning to Earth [87]. The experimental measurements of vBMD in [87] are converted to bone volume fraction assuming that vBMD = 1200 mg HA/cm^3^ corresponds to fully compact bone, *f* = 1. Figure 3a shows that our model calibration is in good agreement with the total vBMD loss experienced by the astronauts in both the tibia, estimated to experience a 60% load reduction in microgravity, and in the radius, estimated to experience a 35% load reduction. Bone recovery after return to Earth (R) is well described by the model in the tibia, but not in the radius, where no significant recovery is observed in the data. The lack of recovery in astronaut radius may be due to irreversible loss of trabecular connectivity, or to the radius serving as a calcium reservoir for increased recovery in the tibia [87, 88]; these effects are not accounted for by our model. Figure 3b compares numerical simulation of the model recalibrated to cortical vBMD loss and recovery from the same experiment [87], which shows that model parameters can be tuned to more closely follow the loss and recovery of bone compartments that have a narrower range of volume fractions. In the rest of the paper, we retain model parameter values corresponding to Figure 3a since these values are derived from our previous mathematical model [34], allowing us to assess the effect of a nonconstant osteocyte density by direct comparison.

**Figure 3.**
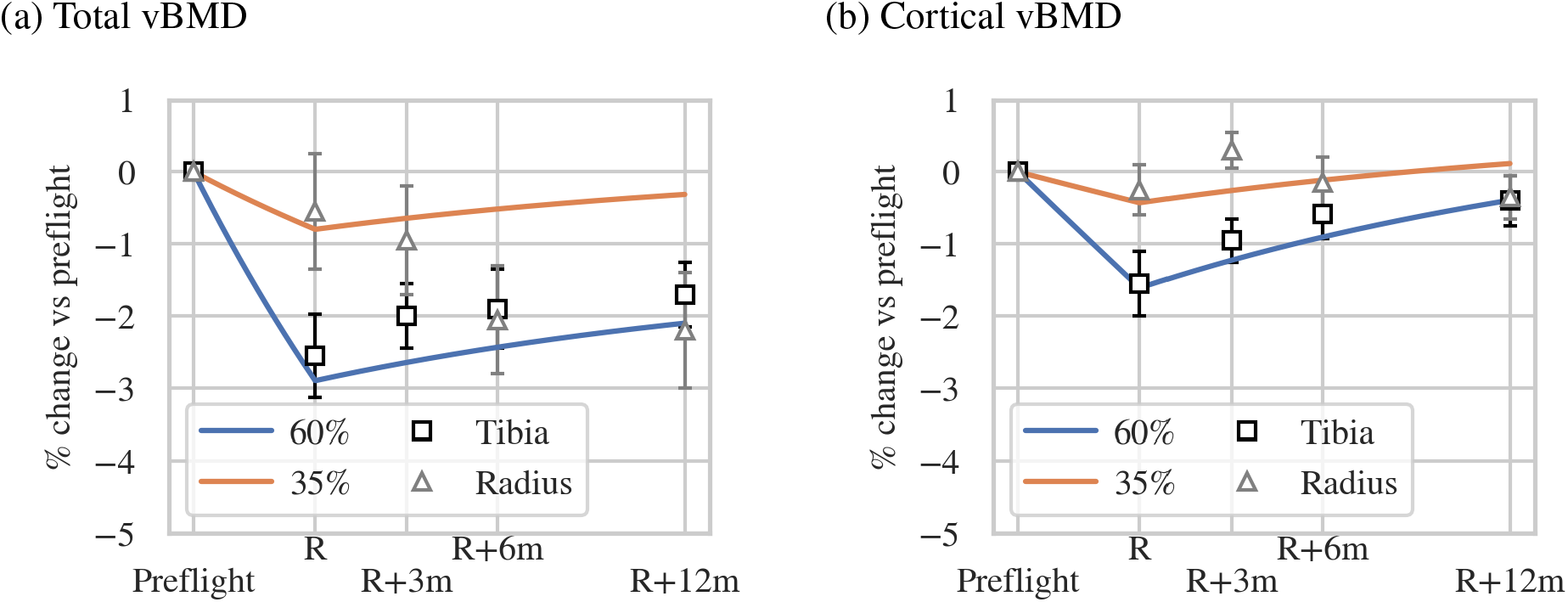
Comparison between long-mission astronaut data from Vico et al. [87] and unloading–reloading model simulations. The label R denotes the time of return to Earth, and corresponds to four months of unloading in the model simulations, after which loads are restored. (a) Numerical simulations of the model assume parameter values listed in Table 2, a preflight bone volume fraction *f* ≈ 0.25 (corresponding to total vBMD ≈ 300 mg HA/cm^3^), and a 60% reduction in loading force in the tibia and 35% reduction in loading force in the radius. The model reproduces the observed loss and recovery of total vBMD in the tibia (blue line) and the loss of total vBMD in the radius (orange line). The lack of recovery in experimental data from the radius may be due to effects not accounted for in our model (see text). (b) The faster recovery rates observed experimentally in cortical vBMD in the tibia and radius can be obtained in the model by increasing the parameter *γ*_Ob_ to *γ*_Ob_ = 2.08*×*10^*−*4^ mm/day. Here simulations assume an initial bone volume fraction *f ≈* 0.75, corresponding to the preflight cortical vBMD *≈* 900 mg HA/cm^3^ measured in the tibia and radius.

### Mechanostat model

In addition to our proposed osteostat model, we present numerical simulations of a more traditional mechanostat model in which there is no change in setpoint, and no change in osteocyte population, i.e., where Eqs (5) and (6) are replaced by a constant setpoint *Ψ*_0_(*t*) = *Ψ*_0_(0) = 3 *×* 10^*−*5^ GPa and a constant osteocyte density 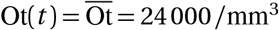, respectively. Other parameters of this mechanostat model are the same as in the osteostat model, except for the value of *γ*_Ob_. The value of *γ*_Ob_ in the mechanostat model is increased by a factor 1.8 (*γ*_Ob_ = 3.75 *×* 10^*−*5^ mm/day), to ensure that the slope of the bone response *f* (*t*) at the onset of reloading matches that of the osteostat model, allowing both models to be contrasted thereafter. The reason why this adjustment is needed is that in the osteostat model, *Ψ*_0_ decreases to a lower value during the unloading phase compared to the mechanostat model.

### 4.2 Long-time unloading and reloading

We start by comparing the evolution of bone in a representative tissue volume TV subjected to long-time unloading and reloading when mechanical adaptation is either (i) a mechanostat with constant setpoint *Ψ*_0_ = const and constant osteocyte density Ot = const; or (ii) our proposed osteostat with a dynamic setpoint adapting according to Eq. (5), and a dynamic population of osteocytes evolving by Eq. (6). The sequence of long-time unloading and reloading that we assume may not correspond to a realistic scenario, but it helps to illustrate key differences in the mechanical response of bone between the osteostat model and the mechanostat model.

As in our previous study [34], the portion of bone is assumed to be initially in mechanical equilibrium at a bone volume fraction of *f* = 0.85, with balanced populations of osteoclasts and osteoblasts. In this initial state, there is no net bone volume loss or gain, but a constant turnover rate 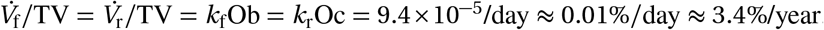, where 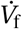 and 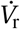 are the volume of bone formed and resorbed in TV, respectively. This turnover value is consistent with values generally assumed in cortical bone [34, 89, 90]. The portion of bone is subjected to a normal loading force *F* =120 N, corresponding to an initial applied macroscopic stress of 30 MPa and initial strain energy density of 3 × 10^*−*5^ GPa [34]. The initial state being in mechanical equilibrium, the setpoint’s initial value is *Ψ*_0_ = 3 × 10^*−*5^ GPa. At time *t* = 0, the loading force is assumed to decrease to either *F* = 78 N (35% decrease), or to *F* = 36 N (70% decrease). After 30 years of unloading, the loading force is restored to its initial value *F* = 120 N.

Figure 4 shows the evolution of several quantities during this long-time unloading and reloading scenario when mechanical adaptation is governed either by the mechanostat (dashed lines), or by the osteostat (solid lines). In both models, at the onset of unloading, the strain energy density *Ψ* drops below the setpoint *Ψ*_0_ (Figure 4e,f), which first induces a mechanics-driven increase in osteoclasts (Figure 4d). The associated decrease in bone volume fraction (Figure 4b) increases bone surface area, which results in a geometry-driven increase in both osteoblasts and osteoclasts (see Eqs (2)–(3)), corresponding to an increased turnover rate with bone loss imbalance (Figure 4c,d). It should be noted here that the increase in osteoblasts is a purely geometrical effect due to the modulation of the populations of osteoblasts and osteoclasts by the specific surface *S*_V_ in Equations (2)–(3). The situation is opposite when unloading from a lower initial bone volume fraction *f* = 0.25. In this case, resorption induced by unloading decreases *S*_V_, and the population of osteoblasts decreases (Supplementary Figure S1).

**Figure 4.**
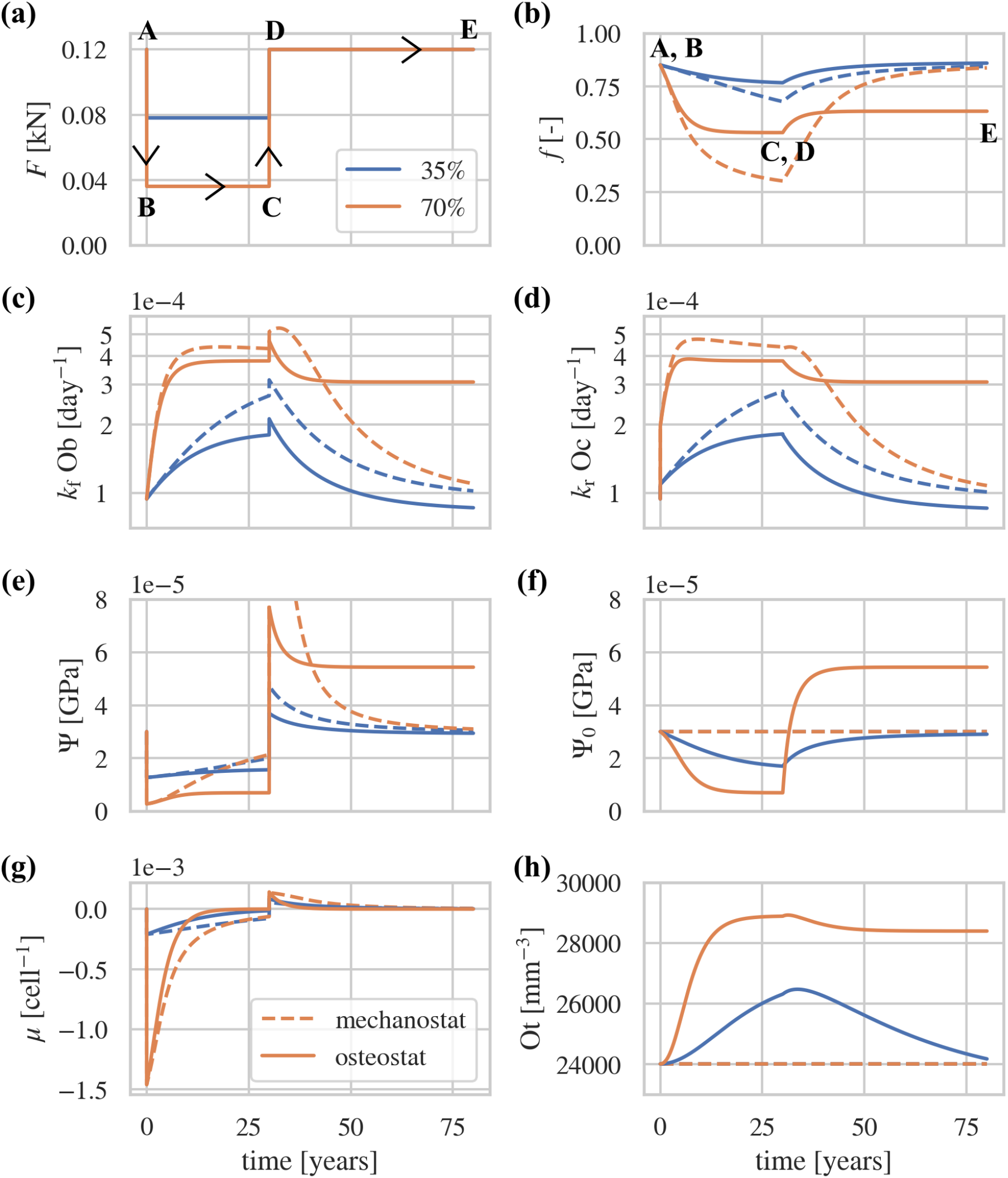
Comparison between mechanostat-based (dashed lines) and osteostat-based mechanical adaptation of bone (solid lines) during long-term unloading and reloading of cortical bone (initial bone volume fraction *f* = 0.85). The plots show the time evolution of (a) the loading force *F* , (b) bone volume fraction *f* , (c) bone formation rate *k*_f_Ob, (d) bone resorption rate *k*_r_Oc, (e) strain energy density *Ψ* , (f) setpoint *Ψ*_0_, (g) mechanical stimulus *µ*, and (h) osteocyte density Ot. Labels A,B denote the transition from normal loads to reduced loads (unloading), and C,D denote the transition from reduced loads back to normal loads (reloading).

When the force is restored to its initial value at *t* = 30 years, the strain energy density *Ψ* jumps above the setpoint *Ψ*_0_, which first induces a mechanics-driven increase in osteoblasts, followed by a recovery of bone volume fraction and a reduction in turnover rate. While these qualitative behaviours of mechanical adaptation are similar in the two models, there are also notable differences, which we now comment upon.

With the mechanostat, the setpoint *Ψ*_0_ is constant (Figure 4f, dashed line), so that the bone volume fraction lost during the unloading phase converges to the percentage reduction in mechanical load at very long times (*f* → 0.255 with 70% unloading, and *f* → 0.5525 with 35% unloading). These values are not quite reached after 30 years of unloading (Figure 4b, dashed lines). Once the loading force reverts to its initial value, all the bone lost is completely recovered; *f* → 0.85 irrespective of the amount of unloading (Figure 4b, dashed lines). The mechanostat model of mechanical adaptation always evolves bone such that its structure is in one-to-one correspondence with its mechanical state. In contrast, with the osteostat, the setpoint *Ψ*_0_ decreases during unloading toward the reduced strain energy density *Ψ* , since new osteocytes generated by remodelling are adapted to lower mechanical loads (Figure 4e, solid lines). The loss of bone during unloading thereby stabilises faster than in the mechanostat and ends up in being less than the percentage reduction in loading (Figure 4b, solid lines). When the loading force returns to its initial value, the recovery of bone depends on how much unloading occurred previously. For a 35% reduction in force, the final bone volume fraction is similar to the initial value, but for a 70% reduction in force, the bone volume fraction after 20 years of reloading (at *t* = 50 years) is only *f* = 0.63 instead of the initial value of 0.85. This is due to the fact that after reloading, the setpoint *Ψ*_0_ adapts to a higher value than its value at time *t* = 0 (Figure 4f, solid orange line). When the force is restored at *t* = 30 years after 70% unloading, the strain energy density *Ψ* jumps to a much higher value compared to after 35% unloading; the density of osteocytes Ot is also higher, and the bone volume fraction *f* is lower. These three effects mean that *Ψ*_0_ converges toward *Ψ* at a much faster rate in Eq. (5). Therefore, *Ψ*_0_ converges to a large value and *f* cannot recover as much bone as has been lost. Contrary to the mechanostat, the osteostat model of mechanical adaptation leads to a loading-history dependence of bone structure, a feature which is observed experimentally [8, 9, 11].

Because of setpoint adaptation, bone volume stabilises to a nonzero value even for a complete removal of the loading force.

Osteocyte density Ot is maintained at a constant value in the mechanostat, but it evolves significantly in the osteostat. The increase of the average density of osteocytes in BV during bone loss (Figure 4h) is due to osteocyte death and the production of new bone by remodelling. Osteocyte death means that older portions of bone have reduced osteocyte density compared to more recent portions of bone. During resorption, the net loss of bone removes portions of bone comprising recent bone, in which lacunar occupancy is high, and old bone, in which lacunar occupancy is low. On average, bone resorption alone does not affect osteocyte density (see Figure 2b). However, the small amount of bone formation that occurs despite net bone loss, replaces a fraction of the bone resorbed, which includes old bone with reduced lacunar occupancy, with new bone in which lacunar occupancy is 100%, which has the effect of increasing the average density of osteocytes within BV. It should be emphasised that the total number of osteocytes within TV, which is proportional to *f* Ot, still decreases with time during this unloading phase. These changes in osteocyte density can also be understood mathematically from Eq. (6). In the initial steady state where dOt*/*d*t* = 0, there are more lacunae than osteocytes. Therefore, Lc_f_ −Ot *>* 0 in Eq. (6), meaning that there must be a balance between the generation of Ot due to remodelling (positive first term in the right hand side of Eq. (6)) and the loss of Ot due to cell death (negative second term in the right hand side of Eq. (6)). When *f* decreases, the positive remodelling-induced generation of osteocyte density becomes greater than the cell-death-induced loss of osteocytes, so that dOt*/*d*t >* 0, meaning that the density of osteocytes increases. When the loading force is restored, osteocyte density in BV decreases again toward a new steady state as the newly formed osteocytes gradually die.

The long-time unloading–reloading scenario simulated in Figure 4 enables us to investigate effective response curves to mechanics (Wolff’s laws). In Figure 5a, we represent the mechanics-induced osteoblast response *k*_f_Ob^(*m*)^ and mechanics-induced osteoclast response *k*_r_Oc^(*m*)^ as a function of the discrepancy between the mechanical variable and the setpoint, *Ψ* (*t*) − *Ψ*_0_(*t*) ≈ *µ*(*t*). In Figure 5b, we represent these mechanics-induced cell populations in response to *Ψ* (*t*) instead. The bone state evolves in time along these response curves as indicated by the labels A–E, defined at specific time points of the unloading–reloading numerical experiment in Figure 4a. Since the mechanostat has constant setpoint *Ψ*_0_, the response curves to *Ψ* (*t*) − *Ψ*_0_(*t*) or to *Ψ* (*t*) are the same in this model except for a horizontal shift. Strikingly, the response curves to *Ψ* (*t*) in the osteostat model (Figure 5b) exhibit a gap between the initial steady state (label A), the steady state reached after unloading (label C), and the steady state reached after reloading (label E). This gap is due to the adaptation of the setpoint *Ψ*_0_(*t*), and not to an assumed “lazy zone” of bone adaptation [2, 3, 14].

**Figure 5.**
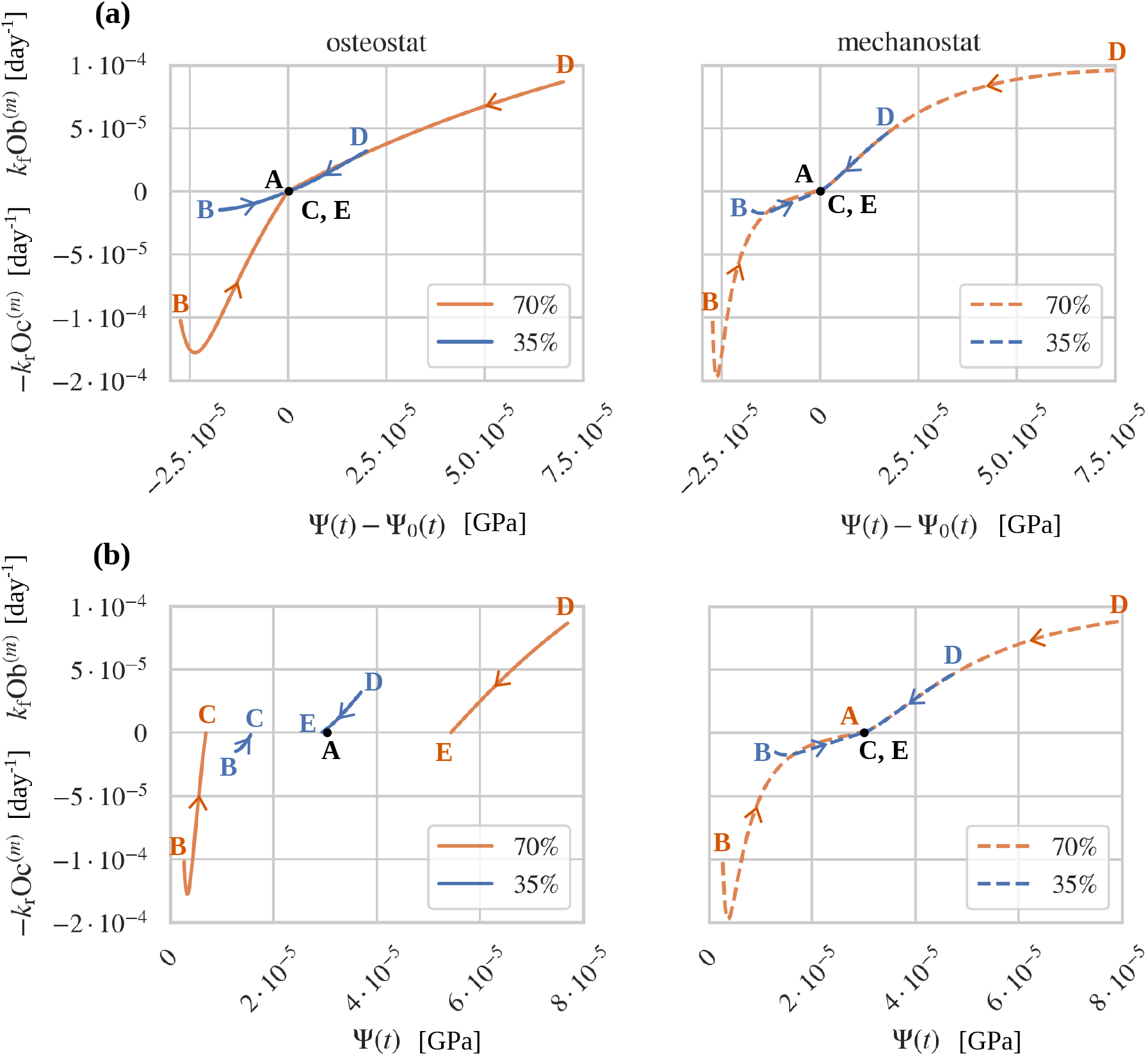
Wolff’s law curves generated with long-term disuse followed by re-loading (see Figure 4). Proposed osteostat with dynamic *ψ*_0_ and Ot (left) and traditional mechanostat with fixed *ψ*_0_ and Ot (right).

Overall, the response curves in Figure 5 in both resorption and formation are such that a greater mechanical stimulus generates a greater response. It should be noted that in early stages of resorption with 70% unloading (label B, orange lines), the osteolytic response reaches a (negative) peak before converging to the new steady-state (label C). This is due to the fact that the initial osteolytic response is first exacerbated by the increase in bone surfaces areas generated by bone resorption. In the mechanostat model, the intensity of force reduction (35% or 70%) generates only slightly different response curves, and only at the start of unloading. In contrast, in the osteostat model, the intensity of force reduction affects the resorption responses at all times because the setpoint adapts.

### 4.3 Mechanical regulation under osteocyte disruptions

Because the osteostat model of mechanical adaptation explicitly accounts for the role of osteocytes in mechanical sensitivity and signalling response, we now explore bone adaptation under osteocyte disruptions. Lacunar occupancy, for example, is known to decrease with age [41, 46], and lacuna density might also reduce in later years of life [38, 41, 45, 74].

A decrease in lacuna occupancy without change in lacuna density can be simulated in our model by increasing the rate of osteocyte apoptosis *A* at a given time, see Eq. (8). Figure 6 shows the time evolution of osteocyte density Ot(*t*) and of bone volume fraction *f* (*t*) when the apoptosis rate *A* is increased by a factor 2 after 5 years of simulation from an initial homeostatic state in mechanical equilibrium with no net bone loss or gain. The increase in osteocyte apoptosis gradually reduces osteocyte density until it converges to a new steady state consistent with the increased value of *A* in Eq. (8) (Figure 6a). Even though osteocyte density decreases, individually, osteocytes do not experience overload or underload. There is no change in mechanics, so the setpoint remains constant. Therefore, there is no mechanical adaptation signal and no mechanics-induced osteoblast or osteoclast generation. The portion of bone continues to be renewed at a constant remodelling rate with no net bone loss or gain (Figure 6b).

**Figure 6.**
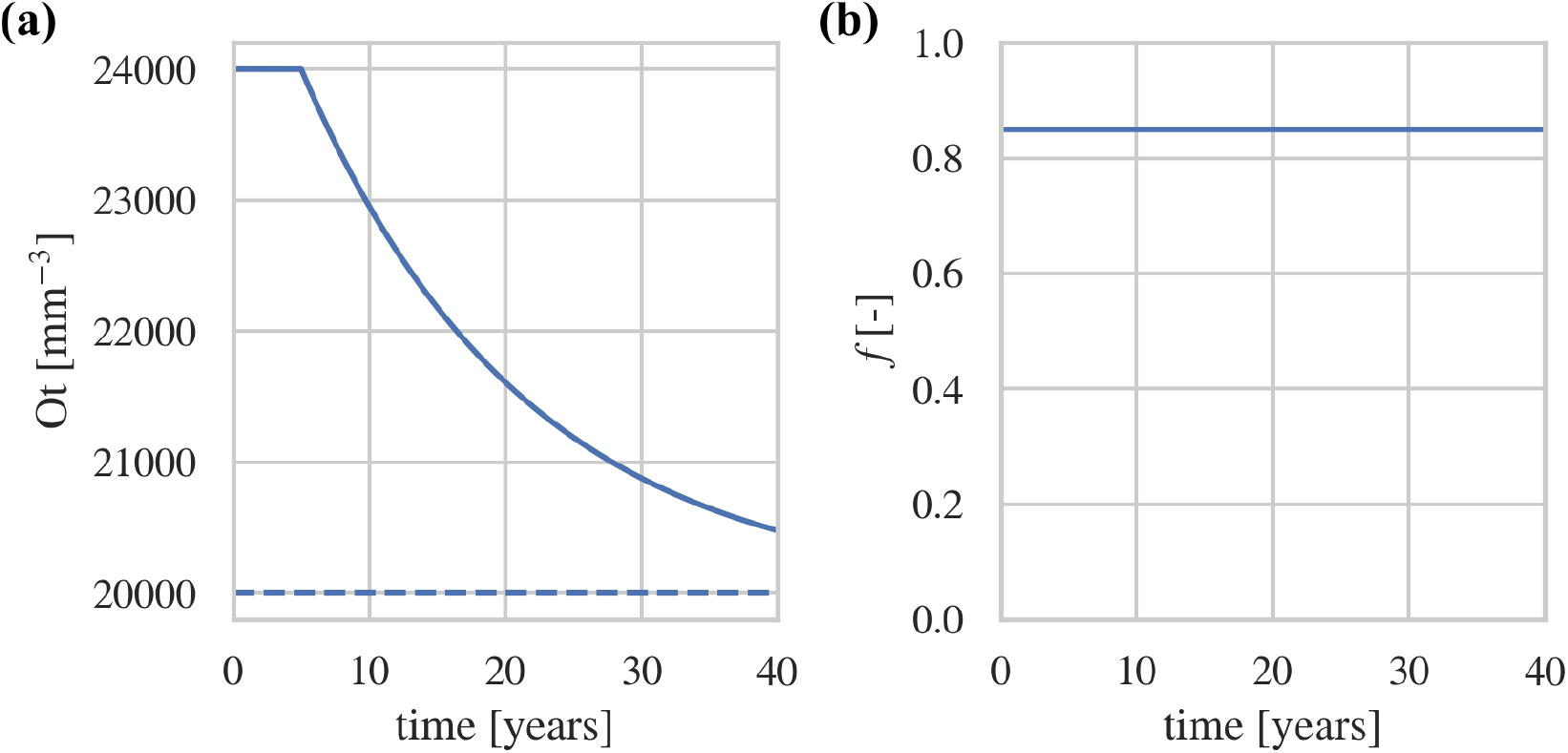
Increasing osteocyte apoptosis *A* by a factor of 2, causes osteocyte density Ot to decrease (a), however, other variables remain unaffected, e.g. *f* (b). The dashed line in (a) represents the new steady-state 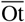 with increased apoptosis *A*.

Changes in osteocyte numbers do not affect mechanical equilibrium, but they affect bone’s mechanical sensitivity and response rate to mechanical loading. Figure 7 compares the evolution of bone volume fraction *f* (*t*) when the portion of bone contains two different initial populations of osteocytes (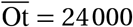and 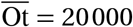) and the loading force *F* is decreased (Figure 7a) or increased (Figure 7b). With lower numbers of osteocytes, the total mechanical signal received by osteoblast and osteoclast precursors in TV is reduced, leading in Eqs (2)–(3) to fewer mechanics-induced osteoclasts (unloading) and fewer mechanics-induced osteoblasts (overloading). The reduction in mechanical sensitivity with lower osteocyte numbers thus results in a reduced amount of bone loss and bone gain, and slower mechanical adaptation responses (Figure 7b). With age, loss of lacuna occupancy could therefore make it more difficult to gain bone, but, according to this model, could also slow down bone loss induced by mechanical disuse. It should be noted here that our osteostat model of mechanical adaptation does not include the presumed role of osteocytes in micro-damage response. It is possible that reduced numbers of osteocytes may lead to more fragile bone tissue due to a lack of detection and removal of micro-cracks. The model also does not account for possible disruptions from osteocyte loss to the baseline of remodelling assumed in the model (*k*_f_Ob^(b)^ and *k*_r_Oc^(b)^), which could affect bone balance even in the absence of mechanical adaptation.

**Figure 7.**
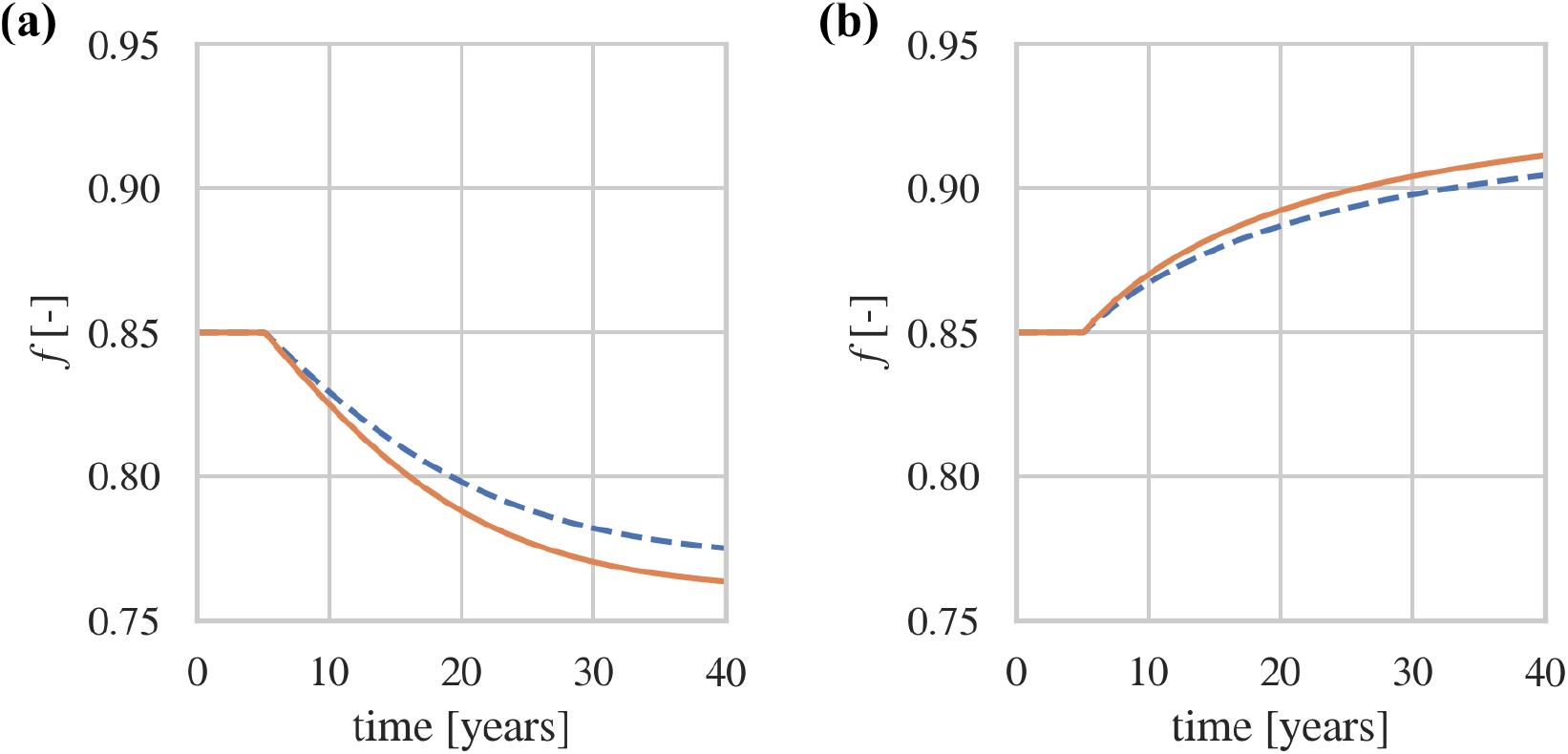
With a change in force, difference in osteocyte density (solid orange: 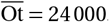, blue dashed: 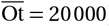) causes slower loss in *f* in disuse (a) as well as a slower gain in *f* in overuse (b).

### 4.4 Mechanical regulation with remodelling imbalance

The regulation of bone tissues is influenced by many factors which may be of biological or mechanical origin. The generation of osteoblasts and osteoclasts involves multiple differentiation stages of precursor cells, each regulated from a combination of signalling molecules and growth factors, that are produced systemically or in response to mechanical signals. Our model accounts for the generation of osteoblasts and osteoclasts due a baseline remodelling rate, attributed to systemic, biological pathways, and osteoblasts and osteoclasts induced by mechanical stimulation. In this section, we consider the bone response regulated by the osteostat model under a biological disruption of remodelling.

Figure 8 shows the time evolution of the bone volume fraction when the baseline remodelling rate is biased toward bone loss, i.e., each bone remodelling unit forms less bone than is resorbed. This is achieved by reducing *α*_Ob_ by 5% compared to *α*_Oc_, to the value *α*_Ob_ = 1.9 *×* 10^*−*4^. This situation could model age-related bone loss, where bone remodelling tends to become more biased toward bone loss due to a lack of bone formation, e.g., due to systemic changes such as in post-menopausal osteoporosis [42,91]. If the loading force is maintained while *α*_Ob_ is reduced, the bone volume fraction decreases due a negative net rate of change d *f /*d*t* (Figure 8, solid blue line). The loss of bone increases mechanical strains and induces a mechanical response that contributes a rate of change of bone volume fraction of d *f* ^(*m*)^*/*d*t* = *k*_f_Ob^(*m*)^ (Figure 8b, dashed line). However, this mechanical compensation is not strong enough to counter the rate of bone loss d *f* ^(*b*)^*/*d*t* = *k*_f_Ob^(*b*)^ − *k*_r_Oc^(*b*)^ induced by the decrease in *α*_Ob_ (Figure 8b, dotted line). As bone loss continues, *ψ* increases further, but because the setpoint *ψ*_0_ gradually adapts toward *ψ*, mechanical stimulation saturates without being able to compensate the biological disruption.

**Figure 8.**
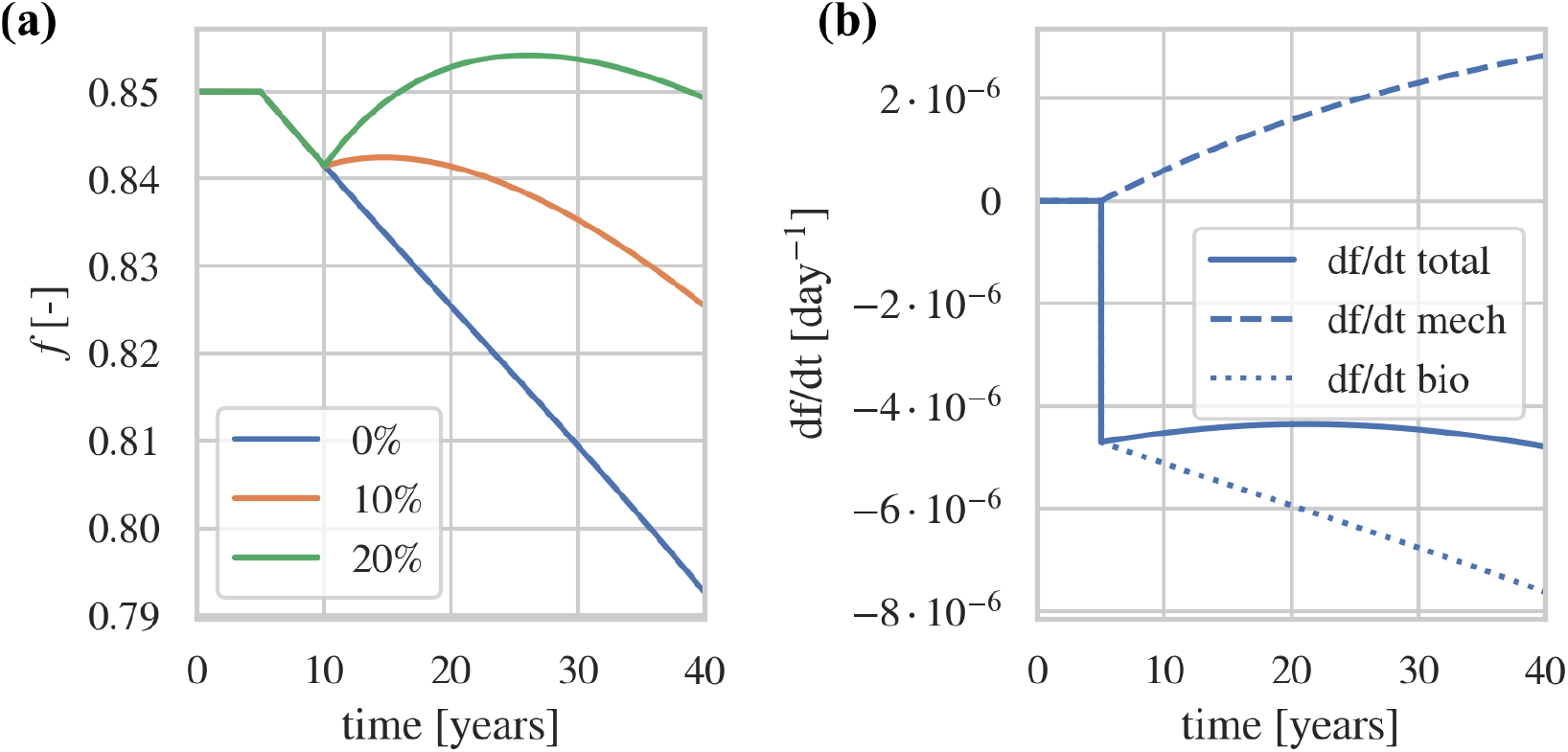
Mechanics-induced compensation of bone loss due to remodelling imbalance. (a) Reducing biochemical formation *α*_Ob_ by 5% (*α*_Ob_ = 1.9 *×* 10^*−*4^) while maintaining the loading force constant results in an approximately constant rate of decrease of *f* (solid blue line). Increasing the loading force by 10% (solid orange line) or 20% (solid green line) can only temporarily help mitigate the loss. (b) The total rate of bone loss for a constant loading force (solid blue line) is the sum of a biological contribution (dotted line) and a mechanics-induced compensation (dashed line) due to bone loss increasing the strain energy density. The mechanics-induced compensation is not strong enough to fully compensate the loss and the net rate of change of bone volume fraction is negative.

Figure 8a also shows the evolution of bone volume fraction when the loading force is increased by 10% (*F* = 123 N, orange line) or 20% (*F* = 148 N, green line) five years after the onset of the biological disruption. The increase in mechanical force curbs the loss and leads to temporary bone gain, but it is not able to fully compensate the biological disruption in the long term.

## 5 Discussion

Experimental observations suggest that osteocyte network morphologies are different in regions of bone experiencing different mechanical loads in homeostasis [30, 31, 61]. These observations support our hypothesis that the mechanical setpoint is encoded in osteocyte properties. The confinement of osteocytes in the lacuno-canalicular network enables osteocytes to sense amplified mechanical stimuli such as fluid flow in the lacuno-canalicular networks [30, 31, 51, 54, 58, 65], and to respond to these stimuli by propagating signals through their extensive network to induce bone formation and bone resorption [22, 28, 47, 92–94]. Different osteocyte properties in different skeletal sites may therefore encode a site-specific mechanical setpoint, while the renewal of osteocytes during bone remodelling may provide a long-term adaptation of the setpoint [34].

The numerical simulations of the mathematical model lead to a number of bone adaptation behaviours that can be put in perspective with experimental observations and previous models of bone adaptation, as we now comment on.

Figures 4 and 5 show that the osteostat theory is similar to the mechanostat theory for small unloadings or short time scales. It is thus compatible with numerous works that have utilised mechanostat theories and Wolff’s type response laws to successfully predict the evolution of bone tissues subjected to mechanical loading [89, 95]. However, the osteostat resolves some of the issues encountered by simple mechanostat theories when there are large unloadings or long time scales. When the setpoint is assumed to be constant (mechanostat model), the percentage of bone loss by unloading corresponds to the percentage of force unloading. In contrast, setpoint adaptation in the osteostat model means that the amount of bone gain or bone loss depends on the loading history, which is observed experimentally [9], rather than being proportional to the net change in loading. This setpoint adaptation prevents total bone loss in regions of bone that experience minimal mechanical loading, such as near the neutral axis of long bones, in the skull, or after spinal cord injury or prolonged bed rest, without needing to include ad-hoc mechanical loading scenarios or load normalisation [89, 96].

With an adapting setpoint, Wolff-type response curves differentiate into two types. The first type is obtained by correlating bone response with the discrepancy between current loads *Ψ* (*t*) and the evolving setpoint *Ψ*_0_(*t*) (Figure 5a). Experimentally, this corresponds to correlating bone responses with an mechano-transduced signal (*µ* in the mathematical model, see Eqs (4), (25)) emitted by osteocytes in response to mechanical stimulation [25, 73, 97]. Potential candidates for such signalling agents are sclerostin, which inhibits osteoblasts [20]; RANKL, which promote osteoclasts [98]; and OPG, which inhibits osteoclasts [99], with these regulatory factors believed to originate from osteocyte intracellular calcium signalling triggered by mechanical stimulation [23].

The second type of Wolff-type response curves are obtained by correlating the bone response with current mechanical loads *Ψ* (*t*) (Figure 5b). Mechanical loads can be estimated in experiments by finite element analysis of the mechanical stresses in bone [16, 19].

For a constant setpoint (mechanostat model), these two types of Wolff response curves have identical shape but are shifted horizontally by *Ψ*_0_. In contrast, the shape of these two types of Wolff response curves differ significantly when the setpoint is adapting (osteostat model), which suggests that this hypothesis may be tested experimentally. For example, by correlating bone formation and resorption responses in an animal loading experiment with either a mechanically transduced signal such as sclerostin [20], or with calculated local mechanical stresses via finite element analysis [16], both type of Wolff response curves may be generated and compared qualitatively to our results.

In our numerical simulations, the second type of Wolff’s response curve in the osteostat model exhibits a response signature that resembles a lazy zone (Figure 5b), but is due to setpoint adaptation. The existence of a lazy zone in Wolff’s response curves, i.e., a region of mechanical stimulus around the mechanical setpoint that does not elicit any mechanics-induced bone response, has been the subject of much debate [14]. While Frost’s mechanostat includes a lazy zone, recent experimental studies measure no lazy zone [16,19]. In our numerical experiments, a steady state with no mechanics-induced bone response can be reached for a range of values of *ψ*. While this seems to be in contradiction to experimental studies, our simulations consider adaptations over long time scales, longer than typical time scales of experimental studies. Additionally, our response curves in Figure 5 isolate the mechanical response, while in experimental studies response curves may include bone formation and bone resorption responses that arise during normal bone remodelling for other reasons than mechanical stimulation. An important distinction of the bone response behaviour in our osteostat model compared to a mechanostat with a lazy zone, is that once the bone has reached a new steady state with a new setpoint, any small change in loading will elicit a small response. In contrast, changes in loading that remain within a true lazy zone would not elicit any bone response.

An interesting consequence of modelling a dynamic osteocyte population is that the average density of osteocytes per BV increases during unloading due to a change in the proportion of old and new bone (Figure 4h). Experimentally, mechanical unloading has been shown to decrease the density of osteocytes, the opposite of what our model shows. However, osteocyte decrease in unloading seems to be related to a mechanically driven increase in osteocyte apoptosis [45, 46, 49]. We did not account for a mechanically driven increase in osteocyte appoptosis in our unloading simulation. This suggests that changes in osteocyte population observed in unloading experiments are the balance between a small increase in osteocyte density induced by an increase in the proportion of new bone by remodelling, and a strong decrease in osteocyte population induced by mechanically driven osteocyte apoptosis.

There is much uncertainty about the lifespan of osteocytes in portions of bone that are seldom remodelled. Osteocytes are believed to be long-living cells more likely to be removed by remodelling than by undergoing apoptosis in a normal homeostatic state [21, 45]. Equation (8) elucidates how the occupancy of lacunae is related to both mechanisms of osteocyte removal: apoptosis (rate parameter *A*), and remodelling (turnover rate *k*_f_Ob). This equation could be used with experimental measurements of osteocyte occupancy to estimate osteocyte lifespan in the absence of remodelling via the rate parameter *A*; the effective half-life of an osteocyte that does not get resorbed is given by ln(2)*/A*. Few studies measure osteocyte occupancy due to the need to measure both filled and empty lacunae [35, 100]. In those that do, the bone volume fraction *f* and the turnover rate of the bone in which these measurements are performed are often not reported. All these quantities are needed in Eq. (8) to estimate *A*.

The onset of age-related bone loss is associated with increased remodelling rate and followed by a net bone volume loss due to a deficit in bone formation in each remodelling event [42, 91]. Mechanical feedback during this process is believed to induce periosteal apposition, but is unable to compensate for endocortical bone loss [42,89]. While our mathematical model does not account for changes in overall bone shape by periosteal apposition, the lack of mechanics-induced compensation of loss of bone volume fraction in TV in Figure 8 is consistent with sustained endocortical bone loss, an effect that may be attributable to the increase in bone surface area as bone volume fraction decreases [79,89,101]. Several exercise programs have been devised to counter age-related bone loss and postmenopausal osteoporosis [102–104]. Our numerical simulations suggest that exercise can indeed help prevent, or even reverse, some of the loss for a significant duration (see Figure 8). However, with reduced osteocyte density in age, exercise interventions might not be as effective (see Figure 7b), as suggested by mechanical unloading and reloading experiments in mice of different ages [105, 106].

Numerical simulations of the mathematical model in Figures 4 and 5 were performed by assuming long-time unloading followed by long-time reloading. The decades time-scale involved in the simulations was chosen to illustrate differences between the osteostat and the mechanostat and to allow the mechanical setpoint to stabilise to a new steady-state, which is slow as the initial state is dense cortical bone (*f* = 0.85) with low remodelling rate. These long time scales also helped to numerically explore the bone response curves in Figure 5. However, long-term changes in mechanical loading do occur after spinal cord injury, prolonged bed rest due to disease or injury, and long space missions. Remodelling rate can accelerate or slow the response of our model to mechanical loading significantly (Eqs (2)–(3)), and it would be valuable to incorporate experimental remodelling rates. Calibrating remodelling rates remains a challenging task for simulation studies. A significant influence on remodelling rate is the availability of bone surface to osteoclasts and osteoblasts, which depends on bone microarchitecture. Some of the nonlinear behaviours of the mathematical model are due to capturing this geometric influence via the specific surface function *S*_V_(*f*) measured experimentally [77–79]. When reported experimentally, bone turnover is often measured using serum concentrations of resporption and formation markers, which are difficult to relate to specific bone sites, and therefore to the bone volume fraction and specific surface [107–109]. Unloading experiments show that cortical and trabecular compartments respond differently to unloading. In particular, bone formation rate is decreased in trabecular bone, similarly to our model when *f* = 0.25 (see Supplementary Figure S1). However, changes to bone formation rate in cortical bone are less clear [88, 110–112]. Some studies suggest increased activation frequency of basic multicellular units with delayed formation [88, 113], which is consistent with the increase in remodelling rate observed in Figure 4, while other studies do not observe statistically significant effects [110].

Our proposed osteostat theory of bone mechanical adaptation has some limitations. There is currently no direct experimental evidence that a mechanical setpoint may be encoded in osteocytes. However, osteocytes and their network have different morphologies in skeletal sites that experience different mechanical loads, which is likely to affect local mechanical sensitivities. Many works have emphasised that the mechanical setpoint is not an inherent mechanical property of matter. It is an effective, emergent property that arises from the many cellular and molecular interactions of a functional syncytium of bone cells [59]. Given the prevalence and location of osteocytes in bone tissue, and their putative role in mechanosensation and mechanoresponse, osteocytes are the most likely candidate for effectively creating a mechanical setpoint, for example through the mechanisms illustrated by our mathematical model of mechanostransduction. Although our hypothesis that the setpoint is embodied in osteocyte properties may be difficult to test directly, the broader implications of an osteostat theory of bone adaptation, which we have explored through mathematical modelling, lead to observable consequences that may provide indirect evidence to validate or invalidate the theory. For example, anosteocytic bone found in teleost fish species, i.e., bone that does not contain osteocytes, is still able to adapt mechanically [114]. There is evidence that osteoblasts and osteoclasts at the bone surface can also sense and respond to mechanical stimulus independently [26]. It would be of high interest to determine whether these types of mechanically responsive anosteocytic bones exhibit mechanical responses that differ qualitatively from those in osteocytic bone, e.g., by being inconsistent with predictions from our osteostat theory.

The mathematical implementation of our osteostat theory of bone adaptation contains further limitations. No spatial dependences are explicitly accounted for in the mathematical model. The evolution of bone structure is assumed to be represented by temporal changes in the bone volume fraction of a fixed region of interest TV of bone tissue. This region is assumed to be large enough to contain multiple simultaneous remodelling units and modelling sites, so that the average density of osteoblasts, osteoclasts, and osteocytes can be described by continuous functions of time. Geometric feedback is included in the model by accounting for specific surface via the phenomenological function *S*_V_(*f*) [77–79, 101]. However, specific surface also depends on bone microarchitecture, such as the prevalence of plates or rods, which is not described by our mathematical model. Similarly, the model does not account for osteocyte network topology and how mechanical signal propagate through the network [30–33]. Osteocyte and lacuno-canalicular network topologies could influence sensing and signalling, and participate in defining the mechanical setpoint. However, the evolution equation (5) for the setpoint does not depend on what osteocytic variables encode the setpoint. It only depends on the assumption that new bone is created in mechanical equilibrium based on current mechanical conditions.

Osteocytes in the mathematical model are assumed to be born fully adapted to the current mechanical loads that prevail in the bone they are created into. This assumption is not particularly supported or contradicted by experimental evidence, but it seems reasonable from a functional perspective and as a working hypothesis in the absence of further information.

The mathematical model also does not include damage of the bone matrix nor bone mineralisation, which both affect mechanics and are dependent on bone remodelling. Osteocytes regulate mineral content [62, 115] and help direct regeneration of damaged bone [45, 46] but the model only includes their role in assisting direct mechanics-induced bone formation and resorption. The lack of sub-TV spatial resolution also means that this temporal model cannot investigate spatio-temporal bone responses to focal damage, nor geometric or mechanical influences on bone formation and resorption that may affect secretory and resorption rates *k*_f_ and *k*_r_ independently of osteoblast and osteoclast populations [116].

Finally, in contrast to our earlier work [34], the model presented here does not account for short time cellular accommodation to mechanics, since we included model features that act on long time scales only, such as osteocyte apoptosis.

## 6 Conclusions

Our proposed osteostat theory makes a number of general assumptions about how bone tissues are regulated biologically and mechanically. These assumptions are reasonably based on experimental observations, and are generally commonly accepted (Section 2). The proposed osteostat model stipulates that the mechanical setpoint of bone is embodied by some properties of the osteocyte network and that these properties may change due to the replacement of osteocytes during remodelling. These properties may involve tethering elements for mechano-sensing, signalling molecules for translating sensed load, and an internal sensitivity. However, this detail does not matter for the broad implications of our osteostat model. The setpoint can be evolved directly under the assumptions that osteocytes are born adapted to the mechanical environment that prevails at the time of their creation. While the general principles of the osteostat are plausible, specific implementation choices of the mathematical model can be expected require refinement as new experimental data becomes available.

The osteostat model highlights many desirable behaviours of bone adaptation that emerge naturally without needing to invoke ad-hoc hypotheses. One major advantage of the osteostat model compared to traditional mechanostat models is that setpoint adaptation prevents full resorption of bone in disuse and unbounded growth under bending [9, 33, 96]. The osteostat model naturally leads to loading history dependent adaptations of bone tissues. While in this work changes to the setpoint were only considered to occur over long time scales due to remodelling, short timescales changes due to cell accommodation [9] can also be included [34]. An interesting outcome of our mathematical model is the possibility to derive other insights. For example, the osteostat yields an equation to estimate the osteocyte half-life due to the possibility of undergoing apoptosis and of being resorbed during remodelling. This equation could be used to estimate osteocyte lifespan by designing new experiments that measure the quantities that it depends on.

When bone response curves are correlated directly with the mechanical variable, the osteostat model exhibits a gap between the osteolytic response and the osteogenic response. This gap may ressemble a lazy zone, but is fundamentally distinct from it. Contrary to a lazy zone, the osteostat model always elicits a bone response. The response curve is shifted horizontally along the mechanical variable axis due to the adapting setpoint, which is what creates a gap. The osteostat model is thereby associated with a family of response curves depending on the current value of the setpoint. The lazy zone has been a long-standing debate [14]. On the one hand, a region of mechanical stimulation where no bone change occurs is necessary for stability when simulating bone adaptation. On the other hand, experimental studies suggest that there is no such zone in either animals or humans [16, 19]. The osteostat model allows us to resolve the discrepancy: there is not necessarily a need to invoke a lazy-zone, bone may always adapt to changes to loading, and stability could be provided by setpoint adaptation.

The precise role of osteocyte disruptions for bone adaptation is an important area of consideration for understanding age-related bone loss. Our model suggests that with less osteocytes, both bone loss and bone gain are slowed, which is partly supported by the literature [117]. Exploring such observable consequences of an osteostat theory further will help refine our understanding of the role of osteocytes in mechanical regulation. Future work will aim to investigate combinations of disruptions to simulate ageing and disease, such as load reduction combined with increased osteocyte apoptosis. While the present work focuses on the evolution of bone volume fraction in a representative tissue volume, there are many opportunities to expand the mathematical model to include spatial interactions and simulate the mechanical adaptation of bone shape and volume in both space and time.

## Acknowledgments

We thank Richard Weinkamer for fruitful discussions. PRB acknowledge support from the Max Planck Queensland Centre on the Materials Science of Extracellular Matrices (MPQC).

## Appendix A Mathematical model

In this appendix, we supplement the mathematical model summarised in Section 3 with additional information and discussion.

### Bone tissue structure

Bone microstructure is expected to be important for turnover rate due to two main factors: (i) the availability of precursor cells in the vascular space of volume fraction 1*− f* , where *f* = BV*/*TV is the bone volume fraction, and (ii) the availability of bone surface for active osteoblasts and osteoclasts to operate on. Due to how bone microarchitecture is organised at different volume fractions, the specific surface

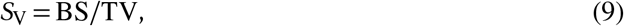

is determined to a great extent by *f* via an experimentally measured relationship *S*_V_(*f*) [77–80]. The relationship *S*_V_(*f*) can be approximated mathematically by

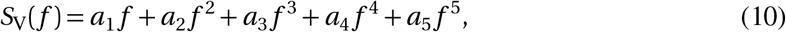

with coefficients *a*_1_ = 14.1*/*mm, *a*_2_ = −10.5*/*mm, *a*_3_ = −17.8*/*mm, *a*_4_ = 43.0*/*mm, *a*_5_ = −28.8*/*mm. These coefficients are slightly adjusted compared to those proposed by Martin [78] to ensure that *S*_V_(0) = *S*_V_(1) = 0 [81] (Figure 9). The relationship *S*_V_(*f*) allows us to represent changes in bone surface area implicity, via changes in bone volume fraction governed by Eq. (1), rather than via modelling the evolution of a three-dimensional bone microstructure. All changes in bone structure induced by formation and resorption are therefore represented in our model by Eq. (1).

**Figure 9.**
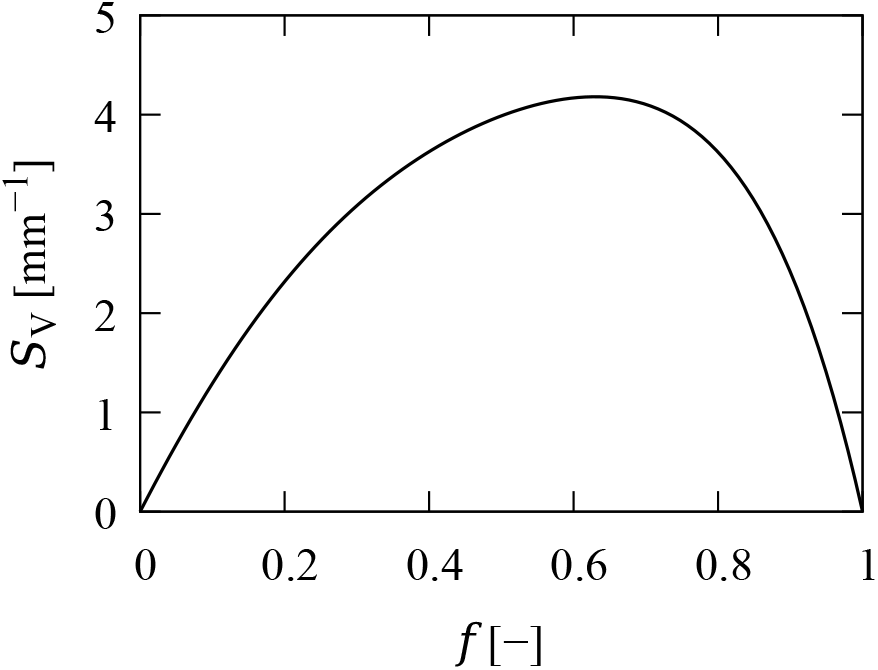
Relationship between specific surface *S*_V_ = BS*/*TV and bone volume fraction BV*/*TV given by Eq. (10), based on experimental measurements [77–81].

### Mechanotransduction and setpoint

Mechanical regulation modifies the populations of osteoblasts and osteoclasts depending on the mechano-transduced signal *µ* (see Eqs (2)–(3)). The signal *µ* represents signalling molecules emitted by osteocytes and received by osteoblasts and osteoclasts. For mathematical convenience, we assume a single, signed transduced signal, such that an osteocyte experiencing overload generates *µ >* 0, and an osteocyte experiencing underload generates *µ <* 0. Whether osteocytes in BV signal an overload response (*µ >* 0) or an underload response (*µ <* 0) depends on (i) current mechanical conditions, and (ii) the sensitivity of osteocytes to these mechanical conditions. Our main assumption is that an osteocyte’s sensitivity to mechanics is set by the mechanical conditions *Ψ* (***r*** , *t*_0_) that prevailed during the generation of the osteocyte at position ***r*** and at time *t*_0_. I.e., the osteocyte’s sensitivity is assumed to be a function of these mechanical conditions, which we define as a mechanical setpoint

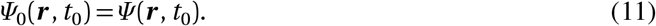

With these assumptions, the mechanical response *µ* of the osteocyte at position ***r*** is zero whenever *Ψ* (***r*** , *t*) = *Ψ*_0_(***r*** , *t*_0_) at later times *t* ⩾ *t*_0_, and nonzero when *Ψ* (***r*** , *t*) ≠ *Ψ*_0_(***r*** , *t*_0_). The mechanical setpoint *Ψ*_0_ at ***r*** remains unchanged until bone is either removed, or re-formed.

To capture these features with a realistic model of osteocyte mechanostransduction, we follow our previous model [34], and consider the osteocyte to be physically attached to the lacuna walls by tethering elements *R* , which transmit mechanical deformation of the lacuna to the osteocyte, and induce a biochemical response in the form of an intracellular signalling compound *S* (Figure 1c). This intracellular compound subsequently induces the generation of an extracellular signal *µ* propagating to the bone surface. The rates at which *S* and *µ* are produced depend on the osteocyte’s sensitivity. Generally, we can conceptualise this model by the sequence of reactions

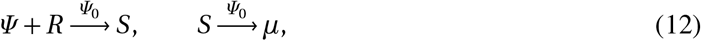

with *Ψ*_0_-dependent reaction rates that represent the osteocyte’s mechanical sensitivity modulating the production rates of *S* and *µ*.

There are several ways by which one may specify these modulations mathematically to ensure that *µ* = 0 whenever *Ψ* = *Ψ*_0_. A simple rate-equation-like model that represents the mechanotransduction (12) in osteocytes may assume a constant number of tethering elements per cell, *R* , no modulation of the generation of *S* by *Ψ*_0_, and a direct (non-differential, fast) modulation of the generation of *µ* by *Ψ*_0_ according to:

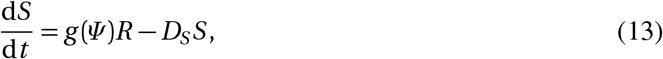

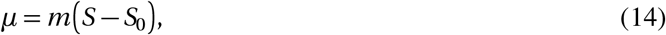

where *m* , *D*_*S*_ , and *R* are constants, *g* (*Ψ*) is a function representing the action of *Ψ* on *R* to generate the signalling compound *S* , and *S*_0_ = *h*(*Ψ*_0_) is a parameter representing the osteocyte’s mechanical sensitivity in generating the signal *µ*. In this model, the setpoint is encoded in the osteocyte-specific parameter *S*_0_. The value of *S*_0_ is set at the osteocyte’s creation time *t*_0_ and position ***r*** by the current mechanical conditions *Ψ*_0_(***r***) = *Ψ* (***r*** , *t*_0_) via the function *h*. Choosing the functions *g* and *h* as 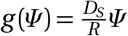 and *h* (*Ψ*_0_) = *Ψ*_0_ for simplicity, one then has, in steady state:

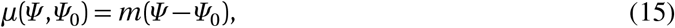

which is the same as Eq. (4). Equation (15) is similar to Wolff’s law, in that it generates an osteogenic response if *Ψ > Ψ*_0_ and an osteolytic response if *Ψ < Ψ*_0_. However, it differs from such a simple response law by proposing a material embodiment of the setpoint *Ψ*_0_ via the osteocyte-specific parameter *S*_0_. This material embodiment of the setpoint allows us to prescribe precisely how *Ψ*_0_ evolves in space and time due to changes in osteocyte properties. The link between the material osteocyte-specific parameter *S*_0_ and the mechanical setpoint *Ψ*_0_ is made explicit by inverting the relationship *S*_0_ = *h*(*Ψ*_0_):

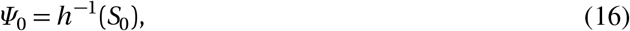

which shows that *S*_0_ embodies a long-term memory of the mechanical strain energy density at which the osteocyte feels neither overloaded nor underloaded. This mechanical setpoint *Ψ*_0_ remains associated with a region of bone until this portion of bone containing the osteocyte is removed by remodelling. Importantly, the mechanical setpoint *Ψ*_0_ associated with a region of bone tissue is not affected by osteocyte death, since osteocytes with new mechanical sensitivies cannot regenerate in empty lacunae. The setpoint *Ψ*_0_ can only change by generating new osteocytes during new bone formation. However, osteocyte death influences the total strength of mechanical response coming from a region of bone tissue, N.Ot *µ* (Figure 2).

The evolution of *S*_0_ due to remodelling is given by setting the value of *S*_0_ in new bone forming at time *t* to *h Ψ* , where *Ψ* represents the current mechanical conditions. Spatially averaging *S*_0_ over BV (the volume referent within which osteocytes are contained), *S*_0_ becomes a function of time only which converges toward the value in newly formed bone, *h* (*Ψ*), at a rate proportional to bone formation rate [34, 83]:

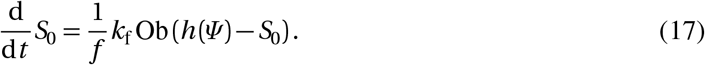

Because the evolution of *S*_0_ in bone is in one-to-one correspondance with the evolution of *Ψ*_0_ via the function *h* , we can also evolve the mechanical setpoint *Ψ*_0_ directly. Any newly formed bone created under mechanical conditions *Ψ* will set the value of *S*_0_ in this portion of bone to *h* (*Ψ*). This corresponds to assigning the setpoint value *Ψ*_0_ associated with this portion of bone to *Ψ* . Similarly to Eq. (17), the evolution of the BV-averaged value of *Ψ*_0_ due to remodelling is thus given by

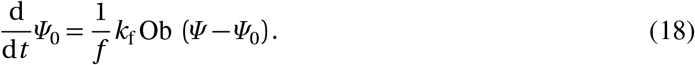

It is clear from Eq. (18) that the detail of how osteocyte-specific properties are encoded in the mechanical setpoint is unimportant. The evolution of the setpoint in BV is entirely determined by the remodelling rate and by what setpoint value is recorded in bone tissue during formation. Here we assume this value to be the current mechanical strain energy density *Ψ* .

In our previous work [34], short-term cell accommodation to mechanical stimulus was included in the mechanotransduction reactions (12) by modulating the number of tethering elements *R* . For all the numerical simulations presented in the current work, we take the long-term behaviour of this more sophisticated model, which leads to a nonlinear function *µ*(*Ψ* , *Ψ*_0_) instead of Eq. (15). This model implements the mechanotransduction reaction sequence in Eq. (12) according to:

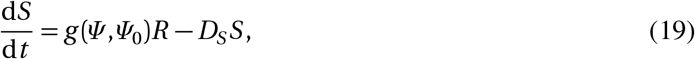

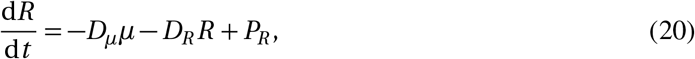

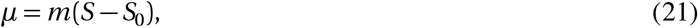

where *D*_*S*_ , *D*_*µ*_, *D*_*R*_ , *P*_*R*_ , and *m* are constant. The term *g* (*Ψ* , *Ψ*_0_)*R* in Eq. (19) captures how the generation of the intracellular signalling compound *S* depends on the number of tethering elements *R* (acting as mechano-receptors), the current mechanical input *Ψ* , and osteocyte sensitivity *Ψ*_0_. The varying number of tethering elements per cell *R* allows osteocytes to partially accommodate to new mechanical loads [9, 34]. Furthermore, the generation of the compound *S* is modulated by *Ψ*_0_, to allow for a saturation of the mechano-transduced response. The function *g* (*Ψ* , *Ψ*_0_) in Eq. (19) is given by

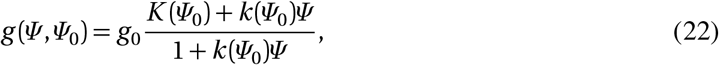

where *K* (*Ψ*_0_) and *k* (*Ψ*_0_) are osteocyte-specific parameters that encode the osteocyte’s mechanical sensitivity through the following explicit expressions:

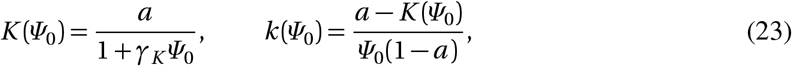

where *a* = *D*_*R*_ *D*_*S*_ *S*_0_*/*(*g*_0_*P*_*R*_), and *γ*_*K*_ is a constant. The expressions in Eqs (22)–(23) and choice of parameter values are based on a number of considerations and constraints on model behaviour, detailed in Ref [34]. Table 2 lists all parameter values used in the model. In particular, these choices are such that if *Ψ* = *Ψ*_0_, then the mechano-transduced signal *µ* in Eq. (21) converges monotonically to 0 in time, after cell desensitisation transients. More precisely, after desensitisation transients, *µ* tends to the long-term value

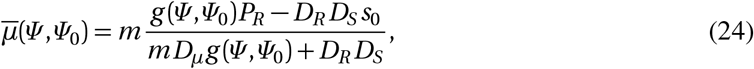

Substituting Eqs (22)–(23) into Eq. (24) provides the following explicit expressions for 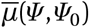:

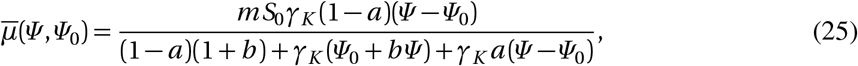

where *b* = *m D*_*µ*_*S*_0_*/P*_*R*_ . In our numerical simulations, we have used the explicit expression in Eq. (25) for the mechano-transduced signal *µ* in Eqs (2)–(3). This nonlinear function still has the property that

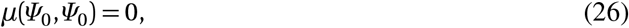

and its linear approximation about *Ψ* = *Ψ*_0_ gives Eq. (4) (up to redefining the numerical value of *m*). The advantage of using the nonlinear function in Eq.(25) is that *µ*(*Ψ* , *Ψ*_0_) saturates to a maximum value when *Ψ* is much greater than *Ψ*_0_, which is sometimes assumed in Wolff-type response curves [3,9,118].

### Osteocytes and osteocyte lacunae

The balance of the population of osteocytes in Eq. (6) is determined by spatially averaging the evolution of the local density Ot(***r*** , *t*) at position ***r*** within BV and time *t* . The derivation of Eq. (6) follows [82, 83] with the only difference being that in the present work, the evolution is coupled with dynamic populations of osteoblasts and osteoclasts. The spatial evolution equation for osteocyte density is

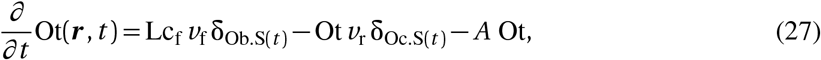

where the first term in the right hand side of Eq. (27) represents the generation of new osteocytes with density Lc_f_ = N.Lc*/*BV at bone-formation surfaces Ob.S(*t*) progressing with speed *v*_f_, the second term represents removal of osteocytes with density Ot at bone-resorption surfaces Oc.S(*t*) progressing with speed *v*_r_, and the third term represents osteocyte cell death. The reader is referred to [83] for full details about how the progression speeds *v*_f_ and *v*_r_ of the forming and resorbing surfaces can be related to *k*_f_Ob and *k*_r_Oc, and how Eq. (27) averages into Eq. (6). These derivations show that if resorption is indiscriminate, i.e., that it does not target bone regions with specific osteocyte densities, the evolution of the BV-averaged osteocyte density Ot does not depend explicitly on bone resorption, as seen in Eq. (6) and illustrated in Figure 2b. It should be emphasised here that the total number of osteocytes within BV is of course affected by resorption.

Lacunar density may itself be time-dependent, since bone formed at different times may contain different lacunar densities. The average lacunar density in BV evolves in time similarly to osteocyte density with the difference that no loss of lacunae (such as resulting from empty lacuna mineralisation) is assumed, other than by resorption:

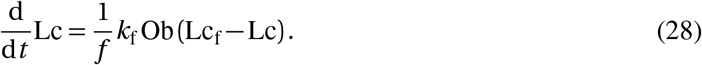

At all times, Ot ⩽ Lc and the average occupancy of lacunae in BV is Ot*/*Lc.

### Osteocyte half-life

The effective half-life of an osteocyte is defined as the time required for the survival probability of an osteocyte to be halved. Since osteocytes may be removed either by bone resorption or by cell death, the probability for an osteocyte to survive to time *t* +*Δt* is the probability it survives to time *t* and that it is not resorbed and does not undergo apoptotis during the time increment

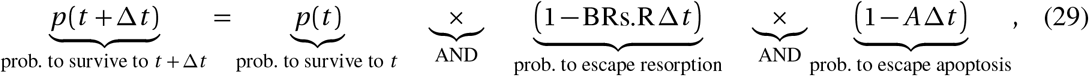

where *A Δt* is the probability for an osteocyte to undergo apoptosis during *Δt* , and BRs.R 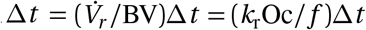 is the probability for an osteocyte to be removed by resorption (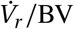 is the fraction of BV removed per unit time). Expanding the products in powers of *Δt* in Eq. (29) gives:

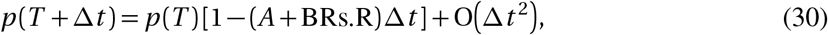

where O (*Δt* ^2^) represent second order powers of *Δt* that become subdominant in the limit where *Δt →* 0. In this limit, using the definition of the time derivative 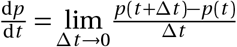, we obtain

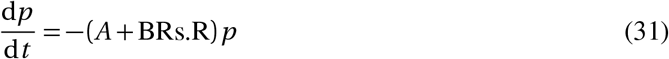

If *A, f* , and BRs.R = *k*_r_Oc = *k*_f_Ob are constant, representing a steady state with a baseline of remodelling rate, then the solution to Eq. (31) is the survival probability function

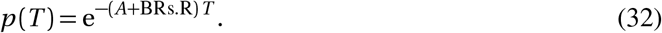

From this function, the half life *T*_1*/*2_ of osteocytes, defined such that *p* (*t* + *T*_1*/*2_) = *p* (*t*)*/*2, is

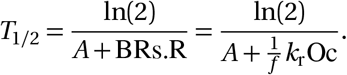

In a dynamic state where *A* or BRs.R are not constant, then *p* (*t*) is still a decreasing function of time, but this function depends on the history of resorption rate and apoptosis rate.

Finally, we note here that steady-state lacuna occupancy in Eq. (8) can also be expressed as:

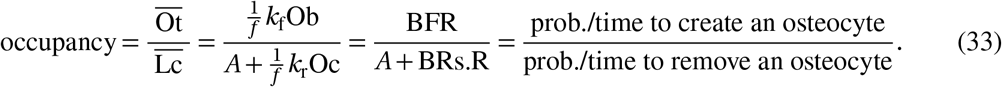

### Strain energy density

Following our previous work [34], we use an estimate of the strain energy density of bone matrix *Ψ* that accounts for stress concentration effects, such that tissue with lower bone volume fraction results in greater strain energy density of bone matrix. This estimate assumes that for small uniaxial deformation, microscopic strains of the bone matrix phase (bm) and of the vascular phase (vas) coincide with tissue-average strains, i.e., *ε*^tissue^ *≈ ε*^bm^ *≈ ε*^vas^, and that tissue-scale stiffness is determined mostly by bone matrix, i.e., *c* ^tissue^ = *f c* ^bm^ + (1 *− f*)*c* ^vas^ *≈ f c* ^bm^. With Hooke’s law at the tissue level, 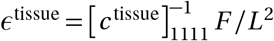, and using 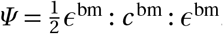, these expressions result in

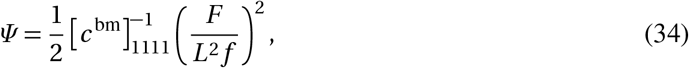

where 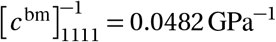, and *F/L*^2^ = 30 MPa. Equation (34) was shown to match numerical evaluations of a micromechanical homogenisation scheme [120] with a maximum relative error of 3% at *f≈* 0, attributed to difficulties of the micromechanical homogenisation scheme at small bone volume fractions [34].

### A.1 Numerical simulations

To solve the presented ordinary differential equations (ODEs) we use Python and the Scipy library. Scipy provides two interfaces to solve ODEs: The original `integrate.ode()` and the newer `ivp_solve()` functions. All experiments were performed using Scipy’s original `integrate.ode()` function with an explicit Runge–Kutta method of order (4)5 (dopri5) [124], asking for values to be calculated every day for long-term unloading reloading experiments and every 1/5 day for all other experiments.

The Scipy ODE solver expects a function that given the current values of the osteostat *f* , Ot and *Ψ*_0_, calculates the corresponding derivatives: d *f /*d*t* using Eq. (1), dOt*/*d*t* using Eq. (6), and d*Ψ*_0_*/*d*t* using Eq. (5).

In addition to the current values for *f* , Ot and *Ψ*_0_, the function receives a vector with force values for each desired time step.

In order to calculate these three derivatives the current force *F* is looked up for the current time step and utilized to calculate *Ψ* from Eq. (34). Next, the value of *Ψ* and the current value of *Ψ*_0_ determine *µ* with the relationship in Eq. (25). Knowing *µ*, the values for *k*_f_Ob and *k*_r_Oc can be calculated using Eq. (2) and Eq. (3), respecively. Note that *µ* is converted to the positive part *µ*^+^ = max(*µ*, 0) *>* 0 or negative part *µ*^*−*^ = *−*min(*µ*, 0) *>* 0. Finally, we obtain d *f /*d*t* using Eq. (1), dOt*/*d*t* using Eq. (6), and d*Ψ*_0_*/*d*t* using Eq. (5).

The implementation of the mechanostat is the same as the osteostat except that *γ*_Ob_ is increased and derivatives for Ot and *Ψ*_0_ were set to 0.0 instead of using Eq. (6) for dOt*/*d*t* and Eq. (5) for d*Ψ*_0_*/*d*t* .

Optionally, the function passed to the ODE solver receives a list of osteostat parameters to change at certain time steps. These parameters change events were used in the experiments in Section 4.3 to increase the apoptosis rate *A* by a factor of 2 after 5 years, and in Section 4.4 to decrease *α*_Ob_ by 5% compared to *α*_Oc_ after 10 years. Changes in force were obtained by changing values in the force vector at the desired time.

Source code for the osteostat, installation instructions and example experiments are available at https://github.com/ypauchard/osteostat.

## Appendix B Evolution of the setpoint and of its encoding parameters

We show in this section that the evolution equation of the setpoint *Ψ*_0_ is independent of the specific model by which we encode *Ψ*_0_ into osteocyte-specific parameters. The evolution of the setpoint only depends on our assumptions that *Ψ*_0_ is set during bone formation at the current value of *Ψ* , and that this value can be reset during bone remodelling.

For illustration, we use the simple mechanotransduction model of Section 3 where *S*_0_ is the osteocyte-specific parameter encoding the setpoint via *Ψ*_0_ = *h*^*−*1^(*S*_0_). It is clear from this relationship that any evolution of *S*_0_ translates into an evolution of *Ψ*_0_. It is important to emphasise that the relationship *S*_0_ = *h* (*Ψ*_0_) is valid at the cellular level, i.e., for the spatio-temporal quantities *S*_0_(***r*** , *t*) and *Ψ*_0_(***r*** , *t*). It is not valid in general for the spatially averaged quantities, unless *h* is linear. To derive the evolution law of the spatially averaged setpoint *Ψ*_0_ in Eq. (5), one may first determine the spatio-temporal evolution of *Ψ*_0_(***r*** , *t*) from that of *S*_0_(***r*** , *t*) via the function *h* , and then proceed to spatial averaging. The evolution of *S*_0_(***r*** , *t*) is governed by a differential equation similar to Eq. (27) [83]:

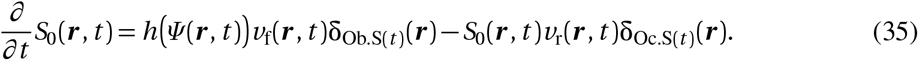

This equation means that at locations ***r*** of bone formation surfaces Ob.S(*t*), the value of *S*_0_ jumps from the value 0 (representing no pre-existing bone) to the value *h* (*Ψ* (***r*** , *t*)) [83]. Similarly, at locations ***r*** of bone resorption surfaces Oc.S(*t*), the value of *S*_0_ jumps from its current value, *S*_0_(***r*** , *t*), to 0. Correspondingly, at locations ***r*** of bone formation surface, the value of the setpoint *Ψ*_0_(***r*** , *t*) = *h*^*−*1^ (*S*_0_(***r*** , *t*)) jumps from the value 0 (no bone) to *h*^*−*1^ (*h Ψ* (***r*** , *t*)))) = *Ψ* (***r*** , *t*), and at locations ***r*** of bone resorption surfaces, the value of *Ψ*_0_ jumps from its current value, to 0. These jumps in *Ψ*_0_ mean that *Ψ*_0_ is equivalently described by the evolution equation:

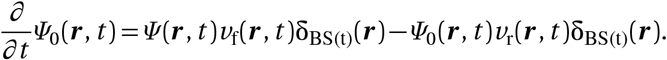

Noticeably, this equation no longer depends on how the setpoint is encoded into osteocyte-specific properties; it is independent of *S*_0_ and *h* . Spatially averaging this equation leads to Eq. (5) [83].

## References

[1] Martin RB (2003) Fatigue damage, remodeling, and the minimization of skeletal weight. J Theor Biol 220:271–276; 10.1006/jtbi.2003.3148

[2] Frost HM (1987) Bone “mass” and the “mechanostat”: a proposal. Anat Rec 219:1–9; 10.1002/ar.1092190104

[3] Frost HM (2003). Bone’s Mechanostat: A 2003 Update. The Anatomical Record Part A: Discoveries in Molecular, Cellular, and Evolutionary Biology 275A:1081–1101; 10.1002/ar.a.10119

[4] Bertram JEA, Swartz SM (1991) The ‘law of bone transformation’: A case of crying Wolff? Biol Rev 66:245–273; 10.1111/j.1469-185X.1991.tb01142.x

[5] Hernandez CJ, Beaupré GS, and Carter DR (2000). “A Model of Mechanobiologic and Metabolic Influences on Bone Adaptation.” Journal of Rehabilitation Research and Development 37:235–244;

[6] Skerry TM (2008) The response of bone to mechanical loading and disuse: Fundamental principles and influences on osteoblast/osteocyte homeostasis. Archives Biochem Biophys 472:117–123; 10.1016/j.abb.2008.02.028

[7] Rosa N et al. (2015) From mechanical stimulus to bone formation: A review. Medical Engineering and Physics 37:719–728; 10.1016/j.medengphy.2015.05.015

[8] Wang C, Fu R, Yang H (2025) Toward a clear relationship between mechanical signals and bone adaptation. Mechanobiology in Medicine 3:100115; 10.1016/j.mbm.2025.100115

[9] Turner CH (1999) Toward a mathematical description of bone biology: The principle of cellular commodation. Calcif Tissue Int 65:466–471; 10.1007/s002239900734

[10] Forwood MR, Turner CH (1995) Skeletal adaptations to mechanical usage: results from tibial loading studies in rats. Bone 17 (4 suppl.):197–205; 10.1016/8756-3282(95)00292-L

[11] Schriefer JL, Warden SJ, Saxon LK, Robling AG, Turner CH (2005) Cellular accommodation and the response of bone to mechanical loading. Journal of Biomechanics 38:1838–1845; 10.1016/j.jbiomech.2004.08.017

[12] Carter DR, Beaupré GS (2001) Skeletal Function and Form (Cambridge University Press); 10.1017/CBO9780511574993

[13] Chen JH, Liu C, You L, Simmons CA (2010) Boning up on Wolff’s law: mechanical regulation of the cells that make and maintain bone. J Biomech 43:108–118; 10.1016/j.jbiomech.2009.09.016

[14] Sugiyama T, Meakin LB, Browne WJ, Galea GL, Price JS, and Lanyon LE (2012) Bones’ Adaptive Response to Mechanical Loading Is Essentially Linear between the Low Strains Associated with Disuse and the High Strains Associated with the Lamellar/Woven Bone Transition. Journal of Bone and Mineral Research 27:1784–1793; 10.1002/jbmr.1599

[15] Meakin LB, Price JS, Lanyon LE (2014) The Contribution of Experimental in vivo Models to Understanding the Mechanisms of Adaptation to Mechanical Loading in Bone. Front Endocrinol 5:154; 10.3389/fendo.2014.00154

[16] Razi H, Birkhold AI, Weinkamer R, Duda GN, Willie BM, Checa S (2015) Aging leads to a dysregulation in mechanically driven bone formation and resorption. J Bone Miner Res 30:1564–1573; 10.1002/jbmr.2528

[17] Gardinier JD, Rostami N, Juliano L, Zhang C (2018) Bone adaptation in response to treadmill exercise in young and adult mice. Bone Reports 8:29–37; 10.1016/j.bonr.2018.01.003

[18] Gardinier JD (2021) The diminishing returns of mechanical loading and potential mechanisms that desensitize osteocytes. Current Osteoporosis Reports 19:436–443;10.1007/s11914-021-00693-9

[19] Christen P, Ito K, Ellouz R, Boutroy S, Sornay-Rendu E, Chapurlat RD, van Rietbergen B (2014) Bone remodelling in humans is load-driven but not lazy. Nat Comm 5:4855; 10.1038/ncomms5855

[20] Robling AG, Niziolek PJ, Baldridge LA, Condon KW, Allen MR, Alam I, Mantila SM, et al. (2008) Mechanical Stimulation of Bone in Vivo Reduces Osteocyte Expression of Sost/Sclerostin. Journal of Biological Chemistry 283:5866–5875; 10.1074/jbc.M705092200

[21] Bonewald LF (2011) The amazing osteocyte. J Bone Miner Res 26:229–238; 10.1002/jbmr.320

[22] Klein-Nulend J, Bakker AD, Bacabac RG, Vatsa A, Weinbaum S (2013) Mechanosensation and transduction in osteocytes. Bone 54:182–190; 10.1016/j.bone.2012.10.013

[23] Morrell AE, Brown GN, Robinson ST, Sattler RL, Baik AD, Zhen G, Cao X, et al. (2018) Mechanically Induced Ca2+ Oscillations in Osteocytes Release Extracellular Vesicles and Enhance Bone Formation. Bone Research 6:6; 10.1038/s41413-018-0007-x

[24] Qin L, Liu W, Cao H, Xiao G (2020) Molecular mechanosensors in osteocytes. Bone Res 8:23; 10.1038/s41413-020-0099-y

[25] Choi JUA, Kijas AW, Lauko J, and Rowan AE (2022) The Mechanosensory Role of Osteocytes and Implications for Bone Health and Disease States. Frontiers in Cell and Developmental Biology 9:770143; 10.3389/fcell.2021.770143

[26] Wang L et al. (2022) Mechanical regulation of bone remodeling. Bone Research 10:16; 10.1038/s41413-022-00190-4

[27] Moriishi T, Komori T (2022) Osteocyte: Their lacunocanalicular structure and mechanoresponses. Int J Molec Sci 23:4373; 10.3390/ijms23084373

[28] Buenzli PR and Sims NA (2015) Quantifying the osteocyte network in the human skeleton. Bone 75:144–150; 10.1016/j.bone.2015.02.016

[29] Mullender MG and Huiskes R (1995) Proposal for the regulatory mechanism of Wolff’s law. J Orthopaed Res 13:503–512; 10.1002/jor.1100130405

[30] Van Tol AF, Repp F, Chen J, Roschger P, Berzlanovich A, Gruber GM, Fratzl P, Weinkamer R (2020) Network architecture strongly influences the fluid flow pattern through the lacunocanalicular network in human osteons. Biomechanics and Modeling in Mechanobiology 19:823–840; 10.1007/s10237-019-01250-1

[31] Van Tol AF et al. (2020) The mechanoresponse of bone is closely related to the osteocyte lacunocanalicular network architecture. Proceedings of the National Academy of Sciences 117:32251–32259; 10.1073/pnas.2011504117

[32] Mehrpooya A, Challis VJ, Buenzli PR (2024) Random walk models for the propagation of signalling molecules in one-dimensional spatial networks and their continuum limit. Proc Roy Soc A 480:20230906; 10.1098/rspa.2023.0906

[33] Mehrpooya A, Challis VJ, Buenzli PR (2026) A mathematical model of osteocyte network control of bone mechanical adaptation. Biomech Model Mechanobiol 52:44; 10.1007/s10237-026-02069-3

[34] Lerebours C & Buenzli PR (2016) Towards a cell-based mechanostat theory of bone: the need to account for osteocyte desensitisation and osteocyte replacement. J Biomech 49:2600–2606; 10.1016/j.jbiomech.2016.05.012

[35] Mullender MG, van der Meer DD, Huiskes R, and Lips P (1996) Osteocyte Density Changes in Aging and Osteoporosis. Bone 18:109–113; 10.1016/8756-3282(95)00444-0

[36] Vashishth D, Verborgt O, Divine G, Schaffler MB, and Fyhrie DP (2000) Decline in Osteocyte Lacunar Density in Human Cortical Bone Is Associated with Accumulation of Microcracks with Age. Bone 26:375–380; 10.1016/S8756-3282(00)00236-2

[37] Qiu S, Rao DS, Palnitkar S, and Parfitt AM (2002) Age and Distance from the Surface but Not Menopause Reduce Osteocyte Density in Human Cancellous Bone. Bone 31:313–318; 10.1016/S8756-3282(02)00819-0

[38] Busse et al. (2010) Decrease in the Osteocyte Lacunar Density Accompanied by Hypermineralized Lacunar Occlusion Reveals Failure and Delay of Remodeling in Aged Human Bone. Aging Cell 9:1065–1075; 10.1111/j.1474-9726.2010.00633.x

[39] Frost HM (1960) In Vivo Osteocyte Death. Journal of Bone and Joint Surgery 42:138;

[40] Dunstan CR, Somers NM, and Evans RA (1993) Osteocyte Death and Hip Fracture. Calcified Tissue International 53:S113–S117; 10.1007/BF01673417

[41] Qiu SD, Rao S, Palnitkar S, and Parfitt AM (2002) Relationships between Osteocyte Density and Bone Formation Rate in Human Cancellous Bone. Bone 31:709–711;10.1016/S8756-3282(02)00907-9

[42] Seeman E (2008) Modeling and remodeling: The cellular machinery responsible for the gain and loss of bone’s material and structural strength. In Bilezikian JP, Raisz LG, Martin TJ, Principles of Bone Biology (3rd Ed) (Academic Press);

[43] Plotkin LI (2014) Apoptotic Osteocytes and the Control of Targeted Bone Resorption. Current Osteoporosis Reports 12:121–126; 10.1007/s11914-014-0194-3

[44] Tiede-Lewis LM, and Dallas SL (2019) Changes in the Osteocyte Lacunocanalicular Network with Aging. Bone 122:101–113; 10.1016/j.bone.2019.01.025

[45] Manolagas SC, Parfitt AM (2013) For whom the bell tolls: Distress signals from long-lived osteocytes and the pathogenesis of metabolic bone diseases. Bone 54:272–278; 10.1016/j.bone.2012.09.017

[46] Jilka RL, Noble B, Weinstein RS (2013) Osteocyte apoptosis. Bone 54:264–271; 10.1016/j.bone.2012.11.038

[47] Cardoso L, Herman BC, Verborgt O, Laudier D, Majeska RJ, Schaffler MB (2009) Osteocyte apoptosis controls activation of intracortical resorption in response to bone fatigue. J Bone Miner Res 24:597–605; 10.1359/jbmr.081210

[48] Gerbaix M, Gnyubkin V, Farlay D, Olivier C, Ammann P, Courbon G, Laroche N, Genthial R, Follet H, Peyrin F, Shenkman B, Gauquelin-Koch G, Vico L (2017) One-month spaceflight compromises the bone microstructure, tissue-level mechanical properties, osteocyte survival and lacunar volume in mature mice skeletons. Sci Rep 7:2659; 10.1038/s41598-017-03014-2

[49] Smith JK (2020) Osteoclasts and microgravity. Life 10:207; 10.3390/life10090207

[50] Ru J-Y, Wang Y-F (2020) Osteocyte apoptosis: the roles and key molecular mechanisms in resorption-related bone diseases. Cell Death and Disease 11:846; 10.1038/s41419-020-03059-8

[51] Weinbaum W, Cowin SC, Zeng Y (1994) A model for the excitation of osteocytes by mechanical loading-induced bone fluid shear stresses. J Biomech 27:339–360; 10.1016/0021-9290(94)90010-8

[52] Turner CH and Forwood MR (1995) What role does the osteocyte network play in bone adaptation? Bone 16:283–285; 10.1016/8756-3282(94)00052-2

[53] Knothe Tate ML, Steck R, Forwood MR, Niederer P (2000) In vivo demonstration of load-induced fluid flow in the rat tibia and its potential implications for processes associated with functional adaptation. The Journal of Experimental Biology 203:2737–2745; 10.1242/jeb.203.18.2737

[54] Fu R and Yang H (2024) Effects of lacunocanalicular morphology and network architecture on fluid dynamic environments of osteocytes and bone mechanoresponses. Physics of Fluids 36:121915; 10.1063/5.0242900

[55] Ganesha T, Laughreya LE, Niroobakhsha M, Lara-Castillob N (2020) Multiscale finite element modeling of mechanical strains and fluid flow in osteocyte lacunocanalicular system. Bone 137:115328; 10.1016/j.bone.2020.115328

[56] Steck R, Knothe Tate ML (2005) In Silico Stochastic Network Models that Emulate the Molecular Sieving Characteristics of Bone. Annals of Biomedical Engineering 33:87–94; 10.1007/s10439-005-8966-7

[57] Adachi T, Kameo Y, Hojo M (2010) Trabecular bone remodelling simulation considering osteocytic response to fluid-induced shear stress. Phil Trans R Soc A 368:2669–2682; 10.1098/rsta.2010.0073

[58] Fu R, Wang C, Shahneela N, Din Ru, Yang H (2025) A whole bone-lacunocanalicular network-osteocyte model examining bone adaptation to distinct loading parameters. Int J Mech Sci 286:109931; 10.1016/j.ijmecsci.2025.109931

[59] Marotti G (2000) The osteocyte as a wiring transmission system. J Musculoskelt Neuron Interact 1:133–136;

[60] Carter Y, Thomas CDL, Clement JG, Peel AG, Hannah K, Cooper DML (2013) Variation in Osteocyte Lacunar Morphology and Density in the Human Femur — a Synchrotron Radiation Micro-CT Study. Bone 52:126–132; 10.1016/j.bone.2012.09.010

[61] van Oers RFM, Wang H, Bacabac RG (2015) Osteocyte shape and mechanical loading. Curr Osteoporosis Rep 13:61–66; 10.1007/s11914-015-0256-1

[62] Kerschnitzki M, Kollmannsberger P, Burghammer M, Duda GN, Weinkamer R, Wagermaier W, Fratzl P (2013) Architecture of the osteocyte network correlates with bone material quality. J Bone Miner Res 28:1837–1845; 10.1002/jbmr.1927

[63] Kollmannsberger P, Kerschnitzki M, Repp F, Wagermaier W, Weinkamer R, Fratzl P (2017) The small world of osteocytes: connectomics of the lacuno-canalicular network in bone. New J Phys 19:073019; 10.1088/1367-2630/aa764b

[64] Chen J, Aido M, Roschger A, van Tol A, Checa S, Willie BM, Weinkamer R (2024) Spatial variations in the osteocyte lacuno-canalicular network density and analysis of the connectomic parameters. PLoS ONE 19:e0303515; 10.1371/journal.pone.0303515

[65] Wang Y, McNamara LM, Schaffler MB, Weinbaum S (2007) A model for the role of integrins in flow induced mechanostransduction in osteocytes. Proc Nat Acad Sci 104:15941–15946; 10.1073/pnas.0707246104

[66] Verbruggen SW, Vaughan TJ, McNamara LM (2014) Fluid flow in the osteocyte mechanical environment: a fluid–structure interaction approach. Biomech Model Mechanobiol 13:85–97; 10.1007/s10237-013-0487-y

[67] Wang H, Du T, Li R, Main RP, Yang H (2022) Interactive effects of various loading parameters on the fluid dynamics within the lacunar-canalicular system for a single osteocyte. Bone 158:116367; 10.1016/j.bone.2022.116367

[68] Martin RB, Burr DB, Sharkey NA (2015) Skeletal Tissue Mechanics (Springer); 10.1007/978-1-4939-3002-9

[69] Franz-Odendaal TA, Hall BK, Witten PE (2006) Buried alive: How osteoblasts become osteocytes. Dev Dyn 235:176–190; 10.1002/dvdy.20603

[70] Bonewald LF, Johnson ML (2008) Osteocytes, mechanosensing and Wnt signaling. Bone 42:606–615; 10.1016/j.bone.2007.12.224

[71] Bonucci E (2009) The osteocyte: the underestimated conductor of the bone orchestra. Rend Fis Acc Lincei 20:237–254; 10.1007/s12210-009-0051-y

[72] Aguirre JI, Plotkin LI, Stewart SA, Weinstein RS, Parfitt AM, Manolagas SC, Bellido T (2006) Osteocyte apoptosis is induced by weightlessness in mice and precedes osteoclast recruitment and bone loss. J Bone Miner Res 21:605–615; 10.1359/JBMR.060107

[73] Schaffler MB, Wing-Yee C, Robert M, and Oran K (2014) Osteocytes: Master Orchestrators of Bone. Calcified Tissue International 94:5–24; 10.1007/s00223-013-9790-y

[74] Milovanovic P, Busse B (2023) Micropetrosis: Osteocyte lacunar mineralization in aging and disease. Current Osteoporosis Reports 21:750–757; 10.1007/s11914-023-00832-4

[75] Manolagas SC (2000) Birth and death of bone cells: basic regulatory mechanisms and implications for the pathogenesis and treatment of osteoporosis. Endocrine Rev 21:115–137; 10.1210/edrv.21.2.0395

[76] Dempster DW, Compston JE, Drezner MK, Glorieux FH, Kanis JA, Malluche H, Meunier PJ, Ott SM, Recker RR, Parfitt AM (2013) Standardized Nomenclature, Symbols, and Units for Bone Histomorphometry: A 2012 Update of the Report of the ASBMR Histomorphometry Nomenclature Committee. J Bone Miner Res 28:1–16; 10.1002/jbmr.1805

[77] Martin RB (1972) The effects of geometric feedback in the development of osteoporosis. J Biomech 5:447–455; 10.1016/0021-9290(72)90003-6

[78] Martin RB (1984) Porosity and specific surface of bone. CRC Critical Reviews in Biomedical Engineering 10:179–222;

[79] Lerebours C, Thomas CDL, Clement JG, Buenzli PR, Pivonka P (2015) The relationship between porosity and specific surface in human cortical bone is subject specific. Bone 72:109– 117; 10.1016/j.bone.2014.11.016

[80] Fyhrie DP, Lang SM, Hoshaw MB, Schaffler MB, Kuo RF (1995) Human vertebral cancellous bone surface distribution. Bone 17:287–291; 10.1016/8756-3282(95)00218-3

[81] Buenzli PR, Thomas CDL, Clement JG, Pivonka P (2013) Endocortical bone loss in osteoporosis: the role of bone surface availability. Int J Numer Meth Biomed Engng 29:1307–1322; 10.1002/cnm.2567

[82] Buenzli PR (2015) Osteocytes as a record of bone formation dynamics: A mathematical model of osteocyte generation in bone matrix. J Theor Biol 364:418–427; 10.1016/j.jtbi.2014.09.028

[83] Buenzli PR (2016) Governing equations of tissue modelling and remodelling: a unified generalised description of surface and bulk balance. PLoS ONE 11:e0152582; 10.1371/journal.pone.0152582

[84] Collet P, Uebelhart D, Vico L, Moro L, Hartmann D, Roth M, Alexandre C (1997) Effects of 1- and 6-month spaceflight on bone mass and biochemistry in two humans. Bone 20:547–551; 10.1016/S8756-3282(97)00052-5

[85] Vico L, Collet P, Guignandon A, Lafage-Proust M-H, Thomas T, Rehailia M, Alexandre C (2000) Effects of long-term microgravity exposure on cancellous and cortical weight-bearing bones of cosmonauts. The Lancet 355:1607–1611; 10.1016/S0140-6736(00)02217-0

[86] Mullender et al. (1996) Osteocyte density changes in aging and osteoporosis. Bone 18:109–113; 10.1016/8756-3282(95)00444-0

[87] Vico L, van Rietbergen B, Vilayphiou N, Linossier M-T, Locrelle H, Normand M, Zouch M, Gerbaix M, Bonnet N, Novikov V, Thomas T, Vassilieva G (2017) Cortical and trabecular bone microstructure did not recover at weight-bearing skeletal sites and progressively deteriorated at non-weight-bearing sites during the year following international space station missions. J Bone Miner Res 32:2010–2021; 10.1002/jbmr.3188

[88] Hargens AR and Vico L (2016) Long-duration bed rest as an analog to microgravity. J Appl Physiol 120:891–903; 10.1152/japplphysiol.00935.2015

[89] Lerebours C, Buenzli PR, Scheiner S, Pivonka P (2016) A multiscale mechanobiological model of bone remodelling predicts site-specific bone loss in the femur during osteoporosis and mechanical disuse. Biomech Model Mechanobiol 15:43–67; 10.1007/s10237-015-0705-x

[90] Parfitt AM (1983) in Bone Histomorphometry, Eds Recker RR;

[91] Seeman E, Martin TJ (2019) Antiresorptive and anabolic agents in the prevention and reversal of bone fragility. Nat Rev Rheumatology 15:225–236; 10.1038/s41584-019-0172-3

[92] van Oers RFM, van Rietbergen B, Ito K, Hilbers PAJ, Huiskes R (2011) A sclerostin-based theory for strain-induced bone formation. Biomech Model Mechanobiol 10:663–670; 10.1007/s10237-010-0264-0

[93] Lewis KJ, Frikha-Benayed D, Louie J, Stephen S, Spray DC, This MM, Seref-Ferlengez Z, Majeska RJ, Weinbaum S, Schaffler MB (2017) Osteocyte calcium signals encode strain magnitude and loading frequency in vivo. Proc Nat Acad Sci 114:11775–11780; 10.1073/pnas.1707863114

[94] Bolamperti S, Villa I, Rubinacci A (2022) Bone remodeling: an operational process ensuring survival and bone mechanical competence. Bone Research 10:48; 10.1038/s41413-022-00219-8

[95] Scheiner S, Pivonka P, Hellmich C (2013) Coupling systems biology with multiscale mechanics, for computer simulations of bone remodeling. Comput Methods Appl Mech Engrg 254:181–196; 10.1016/j.cma.2012.10.015

[96] Levenston ME, Beaupré GS, Carter DR (1998) Loading mode interactions in simulations of long bone cross-sectional adaptation. Comput Meth Biomech Biomed Engng 1:303–319; 10.1080/01495739808936709

[97] Robling AG, Bonewald LF (2020) The Osteocyte: New Insights. Annual Review of Physiology 82:485–506; 10.1146/annurev-physiol-021119-034332

[98] Nakashima T, Hayashi M, Fukunaga T, Kurata K, Oh-hora M, Feng JQ, Bonewald LF, et al. (2011) Evidence for Osteocyte Regulation of Bone Homeostasis through RANKL Expression. Nature Medicine 17:1231–1234; 10.1038/nm.2452

[99] Lacey DL, Boyle WJ, Simonet WS, Kostenuik PJ, Dougall WC, Sullivan JK, San Martin J, Dansey R (2012) Bench to Bedside: Elucidation of the OPG–RANK–RANKL Pathway and the Development of Denosumab. Nature Reviews Drug Discovery 11:401–419; 10.1038/nrd3705

[100] Jordan GR, Loveridge N, Power J, Clarke MT, Parker M, Reeve J (2003) The Ratio of Osteocytic Incorporation to Bone Matrix Formation in Femoral Neck Cancellous Bone: An Enhanced Osteoblast Work Rate in the Vicinity of Hip Osteoarthritis. Calcified Tissue International 72:190–196; 10.1007/s00223-001-2134-3

[101] Buenzli PR, Thomas CDL, Clement JG, Pivonka P (2013) Endocortical bone loss in osteoporosis: The role of bone surface availability. Int J Numer Methods Biomed Eng 29:1307–1322; 10.1002/cnm.2567

[102] Kemmler W, Von Stengel S, Kohl M (2016) Exercise Frequency and Bone Mineral Density Development in Exercising Postmenopausal Osteopenic Women. Is There a Critical Dose of Exercise for Affecting Bone? Results of the Erlangen Fitness and Osteoporosis Prevention Study. Bone 89:1–6; 10.1016/j.bone.2016.04.019

[103] Watson SL, Weeks BK, Weis LJ, Harding AT, Horan SA, Beck BR (2018) High-Intensity Resistance and Impact Training Improves Bone Mineral Density and Physical Function in Postmenopausal Women With Osteopenia and Osteoporosis: The LIFTMOR Randomized Controlled Trial. Journal of Bone and Mineral Research 33:211–220; 10.1002/jbmr.3284

[104] Beck BR (2022) Exercise Prescription for Osteoporosis: Back to Basics. Exercise and Sport Sciences Reviews 50:57–64; 10.1249/JES.0000000000000281

[105] Cunningham HC, West DWD, Baehr LM, Tarke FD, Baar K, Bodine SC, Christiansen BA (2018) Age-dependent bone loss and recovery during hindlimb unloading and subsequent reloading in rats. BMC Musculoskeletal Disorders 19:223; 10.1186/s12891-018-2156-x

[106] Cunningham HC, Orr S Murugesh DK, Hsia AW, Osipov B, Go L, Wu PH, Wong A, Loots GG, Kazakia GJ, Christiansen BA (2023) Differential bone adaptation to mechanical unloading and reloading in young, old, and osteocyte deficient mice. Bone 167:116646; 10.1016/j.bone.2022.116646

[107] Szulc P, Seeman E, Duboeuf F, Sornay-Rendu E, Delmas PD (2006) Bone fragility: failure of periosteal apposition to compensate for increased endocortical resorption in postmenopausal women. J Bone Miner Res 21:1856–1863; 10.1359/jbmr.060904

[108] Burghardt AJ, Kazakia GJ, Sode M, De Papp AE, Link TM, Majumdar S (2010) A Longitudinal HR-pQCT Study of Alendronate Treatment in Postmenopausal Women with Low Bone Density. Journal of Bone and Mineral Research 25:2558–2571; 10.1002/jbmr.157

[109] Malluche HH, Porter DS, Monier-Faugere M-C, Mawad H, Pienkowski D (2012) Differences in Bone Quality in Low- and High-Turnover Renal Osteodystrophy. Journal of the American Society of Nephrology 23:525–532; 10.1681/ASN.2010121253

[110] Zerwekh JE, Ruml LA, Gottschalk F, Pak CYC (1998) The effects of twelve weeks of bed rest on bone histology, biochemical markers of bone turnover, and calcium homeostasis in eleven normal subjects. J Bone Miner Res 13:1594–1601; 10.1359/jbmr.1998.13.10.1594

[111] Bloomfield SA, Allen MR, Hogan HA, Delp MD (2002) Site- and compartment-specific changes in bone with hindlimb unloading in mature adult rats. Bone 31:149–157;10.1016/S8756-3282(02)00785-8

[112] Rolvien T and Amling M (2022) Disuse osteoporosis: Clinical and mechanistic insights. Calcif Tissue Int 10:592–604; 10.1007/s00223-021-00836-1

[113] Minaire P, Meunier P, Edouard C, Bernard J, Courpron P, Bourret J (1974) Quantitative histological data on disuse osteoporosis. Calcif Tiss Res 17:57–73; https://link.springer.com/article/10.1007/BF02547214

[114] Currey JD, Dean MN, Shahar R (2017) Revisiting the links between bone remodelling and osteocytes: Insights from across phyla. Biol Rev 92:1702–1719; 10.1111/brv.12302

[115] Atkins GJ, Findlay DM (2012) Osteocyte regulation of bone mineral: a little give and take. Osteoporosis Int 23:2067–2079; 10.1007/s00198-012-1915-z

[116] Alias MA, Buenzli PR (2018) Osteoblasts infill irregular pores under curvature and porosity controls: a hypothesis-testing analysis of cell behaviours. Biomech Model Mechanobiol 17:1357–1371; 10.1007/s10237-018-1031-x

[117] Tasumi S, Ishii K, Amizuka N, Li M, Kobayashi T, Kohno K, Ito M, Takeshita S, Ikeda K (2007) Targeted ablation of osteocytes induces osteoporosis with defective mechanotransduction. Cell Metabolism 5:464–475; 10.1016/j.cmet.2007.05.001

[118] Wang Y, Wang L, Soro N, Buenzli PR, Li Z, Green N, Tetsworth K, Erbulut D (2024) Bone Ingrowth Simulation Within the Hexanoid, a Novel Scaffold Design. 3D Printing and Additive Manufacturing 11:1949–1960; 10.1089/3dp.2023.0113

[119] Pivonka P, Buenzli PR, Scheiner S, Hellmich C, Dunstan CR (2013) The influence of bone surface availability in bone remodelling — A mathematical model including coupled geometrical and biomechanical regulations of bone cells. Engineering Structures 47:134–147; 10.1016/j.engstruct.2012.09.006

[120] Scheiner S, Pivonka P, Hellmich C (2013) Coupling systems biology with multiscale mechanics, for computer simulations of bone remodeling. Comp Meth App Mech Eng 254:181–196; 10.1016/j.cma.2012.10.015

[121] Grimal Q, Raum K, Gerisch A, Laugier P (2011) A determination of the minimum sizes of representative volume elements for the prediction of cortical bone elastic properties. Biomech Model Mechanobiol 10:925–937; 10.1007/s10237-010-0284-9

[122] Ashman RB, Cowin SC, Van Buskirk WC, Rice JC (1984) A continuous wave technique for the measurement of the elastic properties of cortical bone. J Biomech 17:349–361; 10.1016/0021-9290(84)90029-0

[123] Fritsch A, Hellmich C (2007) ‘Universal’ microstructural patterns in cortical and trabecular, extracellular and extravascular bone materials: micromechanics-based prediction of anisotropic elasticity. J Theor Biol 244:597–620; 10.1016/j.jtbi.2006.09.013

[124] Hairer E, Nørsett SP, Wanner G (1993) Solving Ordinary Differential Equations I, 2nd Ed., Springer Series in Computational Mathematics 8, Springer; 10.1007/978-3-540-78862-1

