## Supplementary Figure S1 for "The “osteostat”: a theory of bone mechanosensing and setpoint adaptation based on osteocytes"

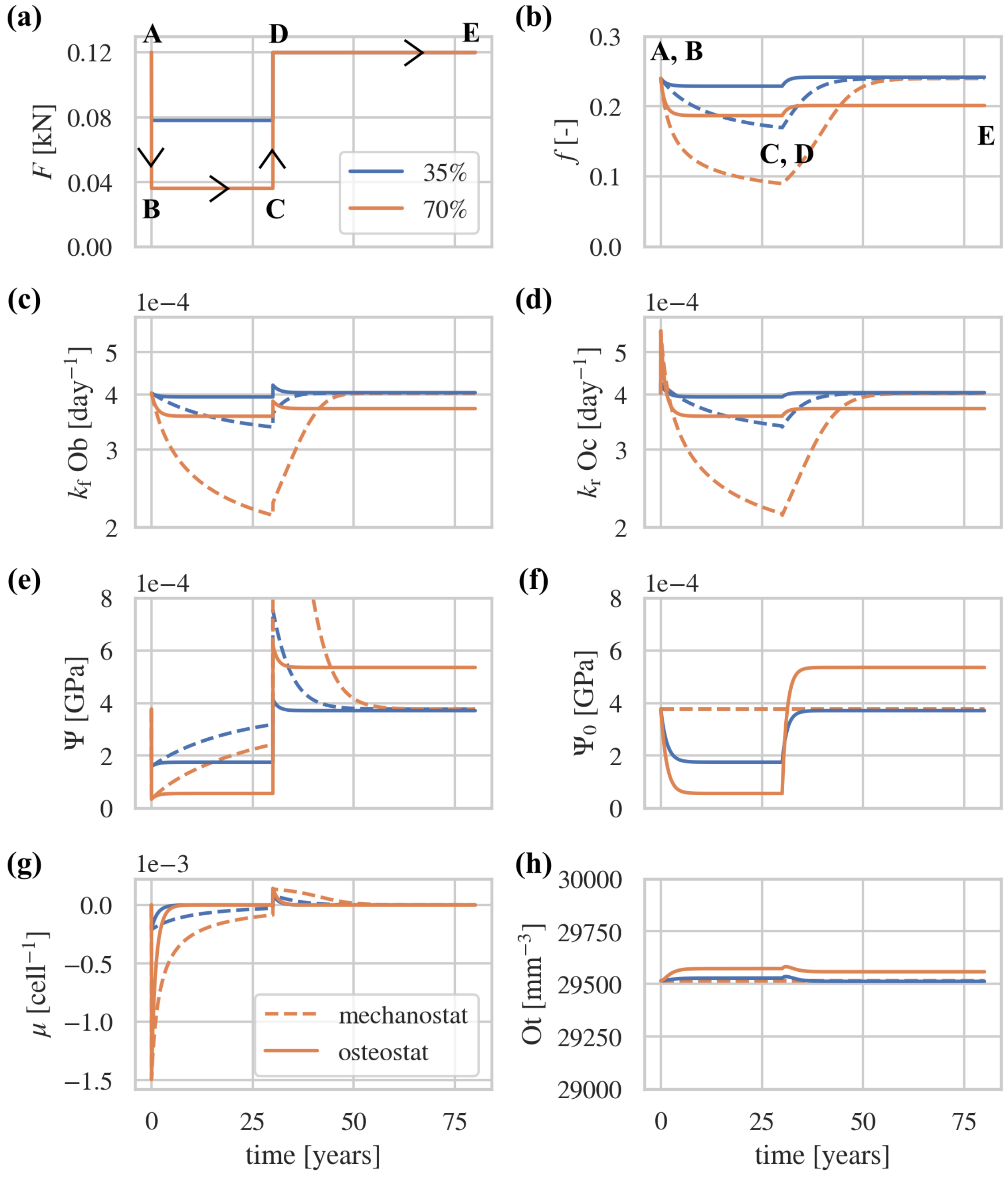


**Figure S1** – Comparison between mechanostat-based (dashed lines) and osteostat-based mechanical adaptation of bone (solid lines) during long-term unloading and reloading of trabecular bone (initial bone volume fraction $f = 0.25$). The plots show the time evolution of (a) the loading force $F$, (b) bone volume fraction $f$, (c) bone formation rate $k_f$Ob, (d) bone resorption rate $k_r$Oc, (e) strain energy density $\Psi$, (f) setpoint $\Psi_0$, (g) mechanical stimulus $\mu$, and (h) osteocyte density Ot. Labels A,B denote the transition from normal loads to reduced loads (unloading), and C,D denote the transition from reduced loads back to normal loads (reloading).
